# Microglial Piezo1 senses Aβ fibrils stiffness to restrict Alzheimer’s disease

**DOI:** 10.1101/2022.03.23.485446

**Authors:** Jin Hu, Hongrui Zhu, Qihua Yang, Huidan Shen, Guolin Chai, Boxin Zhang, Shaoxuan Chen, Qiang Chen, Zhiyu Cai, Xuewen Li, Fan Hong, Hongda Li, Lichao Hou, Wei Mo

**Affiliations:** State Key Laboratory of Cellular Stress Biology, Innovation Center for Cell Signaling Network, School of Life Sciences, Xiamen University, Xiamen, Fujian 361102, China; Department of Anesthesiology, Xiang’an Hospital of Xiamen University, School of Medicine, Xiamen University, Xiamen, China

## Abstract

The pathology of Alzheimer’s disease (AD) is associated with the extracellular amyloid-β (Aβ) plaques that perturb the mechanical properties of brain tissue. Microglia sense and integrate biochemical cues in their local microenvironment, intimate linking with AD progress. However, neither the microglial mechanosensing pathway nor its impact on AD pathogenesis is well studied. Here, we showed that the mechanosensitive ion channel Piezo1 is increased in microglia upon stiffness stimuli of Aβ fibrils. The upregulation of Piezo1 in Aβ plaque-associated microglia was observed in AD mouse models and human patients. Microglia lacking *Piezo1* disturbed microglial clustering, phagocytosis, and compaction of Aβ plaques, resulting in the exacerbation of Aβ and neurodegenerative pathologies in AD. Conversely, pharmacological activation of Piezo1 ameliorated brain Aβ burden and cognitive impairment in the 5×FAD mouse. Together, our results reveal Piezo1, as a mechanosensor of Aβ fibrils stiffness in microglia, could represent a promising therapeutic target for AD.

## Introduction

Alzheimer’s disease (AD) is the most common progressive neurodegenerative dementia in the elderly. Pathologically, AD is characterized by the presence of extracellular amyloid-β (Aβ) plaques and intracellular neurofibrillary tangles containing phosphorylated tau protein, both linked to neuronal damage, synapse dysfunction, and subsequent cognitive impairments (Bloom, 2014; Pereira et al., 2021). The preponderance of human genetics evidence has shed light on a number of common and rare genetic loci closely associated with AD risk. Notably, many of these risk loci are associated with genes that have been reported as microglia specific or highly expressed in microglia relative to other cell types in the central nervous system (CNS) (Lopes et al., 2022; Verheijen and Sleegers, 2018). This broad trend implicates microglia, as the predominant immune cells residing in the brain parenchymal have emerged as crucial players in the pathogenesis of AD.

The diversity of microglial functions results from their ability to respond dynamically to the local microenvironmental cues (Li and Barres, 2018). Microglia actively recognize and phagocytose Aβ aggregates tightly wrap around compact plaques and contribute a leak-tight physical barrier that avoids the outward extension of amyloid fibrils (Condello et al., 2015; Hansen et al., 2018). The complex chemical components of Aβ plaque, such as misfolded proteins, nucleic acids, and lipids, trigger microglia responses via their specific receptors (Drummond et al., 2017; Stewart and Radford, 2017). Previously study reported that Aβ plaques possess a rigid core formed by densely folded Aβ sheets (Mattana et al., 2017). Physical signals, including stiffness or roughness of matrix architecture, can equally enhance immune cell recruitment and activation (Ayata and Schaefer, 2020; Jain et al., 2019; Upadhyaya, 2017). Following the distinct physical properties of the Aβ plaques, it is tempting to speculate that Aβ plaques present a mechanical stimulus to microglia, and they may act as mechanosensing cells in the pathogenesis of AD. Still, how microglia sense and respond to their mechanical microenvironment changes in the development of various neurodegenerative pathological states, such as Aβ deposition in AD, have not been elucidated as yet.

Mechanical force induces cellular functions entails mechanosensitive proteins, known as mechanosensory ion channels (MSICs)(Delmas et al., 2011). Microglia express several MSICs, which process microglial activation, trophic support, and pain, mainly by modulating intracellular calcium (Ca^2+^) homeostasis (Echeverry et al., 2016). Piezo1 is evolutionarily conserved and activated by various mechanical stimuli, including shear stress, stiffness, and cyclical pressure across multiple cell types, contributing to Ca^2+^ signaling via permeating extracellular Ca^2+^ influx (Atcha et al., 2021; Coste et al., 2010; Ranade et al., 2014; Solis et al., 2019). A significant increase of Piezo1 at protein aspects has been observed in the Aβ-deposited hippocampus in mice (Kim et al., 2019). However, whether Piezo1 contributes to microglia mechanosensing of Aβ plaques, either in beneficial or detrimental ways that affect AD’s progression, has not been explored yet.

Here, we discover for the first time the disparity between the mechanical properties of the Aβ plaque-associated brain tissues (PAT) and non-Aβ plaque-associated brain tissues (NPAT) far away from the core of amyloid plaques in live 5×FAD mouse brain slices. Next, we identify the Piezo1 ion channel as a classical MSICs in microglia, which sustains microglial response to Aβ fibrils stiffness. Moreover, microglial Piezo1 upregulation is proven to be strictly Aβ plaque-associated in 5×FAD mice and AD patients. Noticeably, we discern that microglial *Piezo1* deficiency impairs microglia surrounding, phagocytosing, and compacting Aβ plaques, thereby promoting the further deterioration in 5×FAD mice, suggesting the critical role of microglial Piezo1 in preventing or controlling the progression of Aβ pathology in AD. Subsequently, systemic administration of the Piezo1 specific agonist, Yoda1, reduces Aβ burden and ameliorates cognitive decline in 5×FAD mice, indicating Piezo1 may therefore be a potential therapeutic target for the treatment of AD.

## Results

### Microglia responses to the stiffness of β-amyloid (Aβ) fibrils/insoluble aggregates via Piezo1

Aβ plaques in the brain of AD are mainly composed of the insoluble Aβ fibrils, which reveal a remarkable stiffness (Mattana et al., 2017). However, the stiffness discrepancies between Aβ plaque-associated tissues (PAT) and non-Aβ plaque-associated tissues (NPAT) in the brain have not been identified adequately. To measure the stiffness of the PAT, we took advantage of *in vivo* Aβ plaque labeling using methoxy-X04 (MX04) and performed atomic force microscopy (AFM) indentation experiments on uninjured 5×FAD mouse brain slices perfused with artificial cerebrospinal fluid (Figure 1A). As previously stated, Hertzian contact theory is used to calculate the elastic modulus of soft materials, including brain tissues (Moeendarbary et al., 2017). In our system, the force indentation curves were consistent with the slope of the Hertz contact model (Figure 1B). Elastic modulus values indicated that MX04^+^ PAT with the apparent elastic modulus of 582.8 ± 109.9 (Mean ± SD) Pa, while the MX04^−^ NPAT was 317.9 ± 33.78 Pa, suggesting a significantly higher stiffness of PAT than NPAT (Figure 1C), which may be attributable to the stiff Aβ aggregates in the plaques.

**Figure 1.**
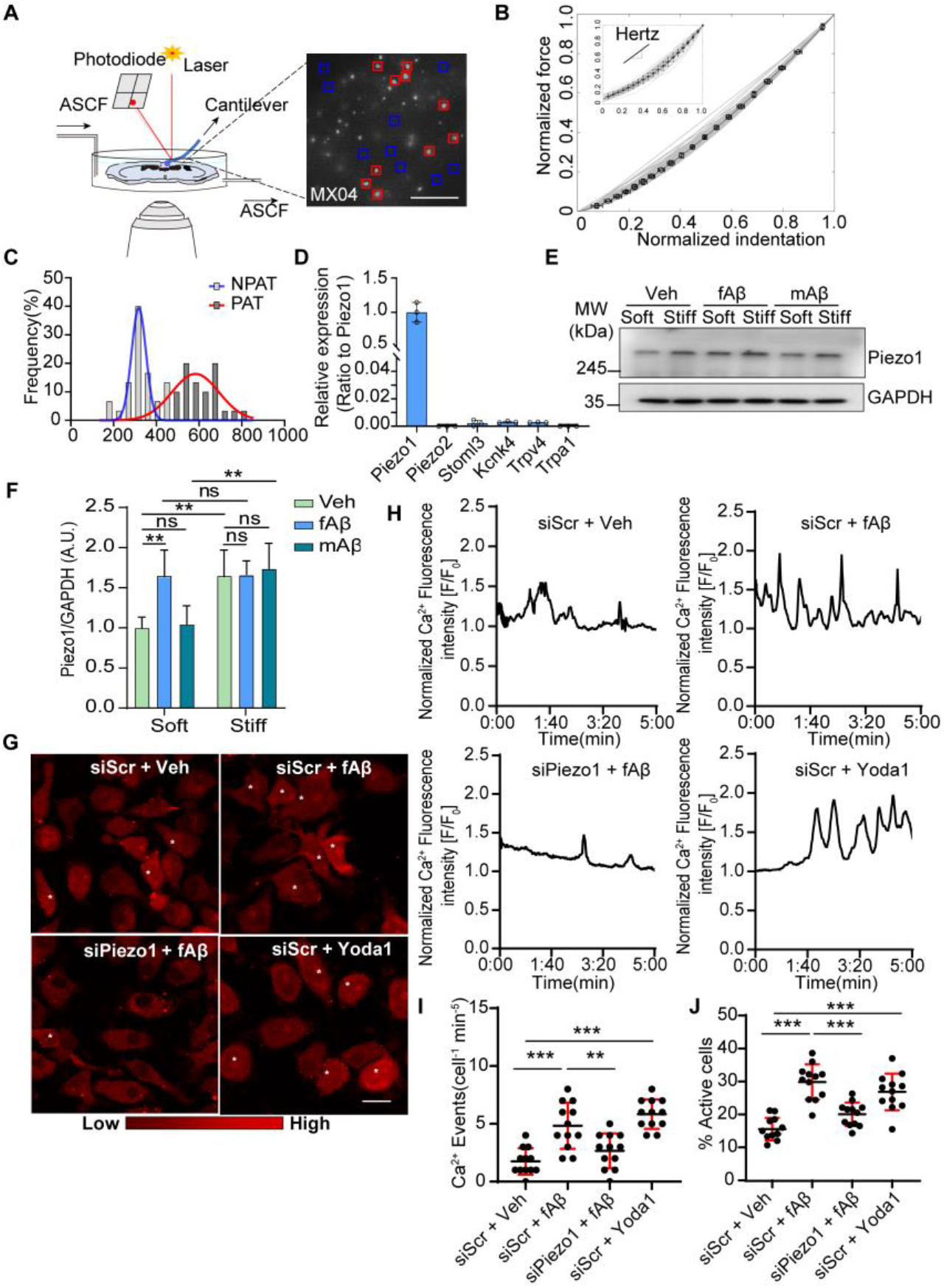
Aβ Fibrils Stiffness Sensing Requires Piezo1 in Microglia. (A) Schematic drawing of AFM measurements of cultured brain slice. MX04 positive Aβ plaque-associated tissues (PAT) and MX04 negative non-Aβ plaque-associated tissues (NPAT) were labeled with red and blue squares. Scale bar, 300 μm. (B) The fitting force–indentation curves were calculated with a spherical Hertzian contact model according to each force–indention curve. Normalized idendation was obtained from the vertical cantilever tip position, and normalized force was calculated from vertical deflection according to every force–indentation curve. (C) Frequency distribution of Young’s Modulus (Pa) according to the outcomes from PAT and NPAT measured by AFM. (D) RT–qPCR analysis of known mammalian mechanosensory ion channels from the primary microglia (n=3 independent experiments)). (E and F) Immunoblotting and statistical analysis of the Piezo1 protein levels in primary microglia on soft or stiff hydrogel stimulated with fAβ (2 μM) or mAβ (2 μM) for 4 h (n=4 independent experiments). (G) Representative images reflect traces of individual Ca^2+^ events of primary microglia seeded on the soft gel (Asterisks denote the occurrence of a Ca^2+^ event). Scale bar, 20 μm. (H) Dynamic traces of individual Ca^2+^ events according to normalized real-time fluorescence Intensity (F/F_0_). (I) The number of Ca^2+^ events in each primary microglia during 5 min observation (n = 12 independent view fields per group). (J) The percentage of flickering cells in a single high-power field (n = 12 independent view fields per group). Statistics: Mean ± SEM. two-way ANOVA with Bonferonni post hoc analyses(F); one-way ANOVA with Bonferroni post hoc test (I and J); ns, no significance; *p < 0.05; **p < 0.01; ***p < 0.005.

Environmental stiffness correlates with mechanosensation-mediated cell functions, including migration, differentiation, and immune response (Chakraborty et al., 2021; Nourse and Pathak, 2017; Segel et al., 2019). Microglia are versatile effectors under CNS physiological and pathological conditions (Hanisch and Kettenmann, 2007) and display a preference toward stiff materials, which is called durotaxis (Bollmann et al., 2015). We investigated several well-studied MSICs expressions in microglia from published datasets from mouse (GSE52564) to human (GSE99074) (Galatro et al., 2017; Zhang et al., 2014) and confirmed the highest expression of Piezo1 compared to other well-known MSICs in microglia (Figures 1D, Figures S1A-S1B). Although the Piezo1 channel has been reported as a mechanical sensor in many cell types (Murthy et al., 2017), its role in microglial stiffness sensing is unclear. To investigate whether microglial Piezo1 directly responds to Aβ fibrils stiffness, soluble Aβ_1–42_ monomers (mAβ42) and insoluble Aβ_1–42_ fibrils (fAβ42) samples were prepared for subsequent experiments (Figure S1C). As expected, fAβ42 exhibited high rigidity (Median of 95.84 MPa), as detected by AFM (Figures S1D-S1E). Polyacrylamide-based hydrogels are now widely used in cell-substrates mechanical interactions (Kandow et al., 2007; Tse and Engler, 2010). We then evaluated the effects of fAβ42 and mAβ42 treatment on the expression of Piezo1 in primary microglia cultured on soft (546.5 ± 43.9 Pa) and stiff (11.03 ± 0.58 MPa) polyacrylamide hydrogels, respectively (Figure S1F). The mRNA level of Piezo1 was almost the same in the entire experiment (Figure S1G). In contrast, Piezo1 protein was increased in microglia upon stiff stimulation (stiff hydrogel vs. soft hydrogel (Figure 1E, line 2 vs. line1); fAβ42 vs. mAβ42 (Figure 1E, line 3 vs. line5 and Figure 1F, soft panel), suggesting microglia sense the stiffness of Aβ via Peizo1. When stiffness perception was saturated in microglia with stiff hydrogel culture, neither fAβ42 nor mAβ42 treatment further induced the upregulation of Peizo1 (Figure 1E, line 4 and 6 vs. line 2 and Figure 1F, stiff panel), indicating the fAβ42-induced elevation of piezo1 protein level in microglia was predominantly stimulated by fAβ42 stiffness but not by fAβ42 chemistry.

Piezo1 is a nonselective ion channel that exhibits a preference for Ca^2+^ in response to mechanical stimuli (Coste et al., 2010; Gnanasambandam et al., 2015). We next examined whether Piezo1 regulates Ca^2+^ activity in response to fAβ42 mediated microglial stiffness sensing. We found that Piezo1 knock-down by short interfering RNA (siRNA) (Figures S1H-S1J) significantly abolished Ca^2+^ activity due to fAβ42 stimulation in primary microglia cultured on the soft hydrogel, as shown by impeding the increase of single-cell Ca^2+^ events and reducing the percentage of active cells involved in Ca^2+^ influx (Figures 1G-1I). Together, these results implicate that the stiffness of fAβ42 as a physical cue drives mechanosensation-mediated signal transduction via microglial Piezo1.

### The upregulation of microglial Piezo1 is disease-associated

To examine whether Aβ plaque stiffness induces Piezo1 expression in microglia *in vivo*, we detected the expression pattern of Piezo1 after stereotactic FAM-labeled Aβ42 fibrils (FAM-fAβ42) injection in the hippocampus of *Piezo1^P1tdT^* (endogenous Piezo1 protein fused with tdTomato) (Ranade et al., 2014) mouse brain. We found strong signals from Piezo1-tdTomato in the proximity of the FAM-fAβ42 deposition, as shown by the RFP staining (Figure S2A). Similarly, Iba1 positive microglia contacting FAM-fAβ42 deposition displayed higher Piezo1 expression than those not touching FAM-fAβ42 deposition (Figure S2B). To investigate whether the upregulation of microglial Peizo1 is disease-associated, we examine the expression of Peizo1 during the formation of Aβ plaque that is closely related to AD pathogenesis. 5×FAD mice are mouse model of AD with rapid amyloid pathology. We crossed *Piezo1*^*P1-tdT*^ mice with 5×FAD mice to create 5×FAD mice that expressed Piezo1-tdTomato. Piezo1 expression was visualized by immunofluorescent staining with anti-RFP antibody and found on microglia positive for Iba1 (Figure 2A). Quantification of Piezo1-tdTomato fluorescence indicated that robust upregulation of Piezo1 protein occurred in plaque-associated microglia (PAM, 582.8 ± 109.9 Pa) versus those non-plaque associated microglia (NPAM, 317.9 ± 33.78 Pa) (Figure 2B). Finally, the clinical relevance comes from the analysis of postmortem brain tissue from AD patients (Braak stages IV) (Figure S2C) revealed that PIEZO1-enrichment microglia predominantly congregated around Aβ plaques (Figures 2C-2D), consistent with the results obtained in the AD mouse model. In summary, these results indicate that microglial PIEZO1 is specifically up-regulated on and around Aβ plaques in AD.

**Figure 2.**
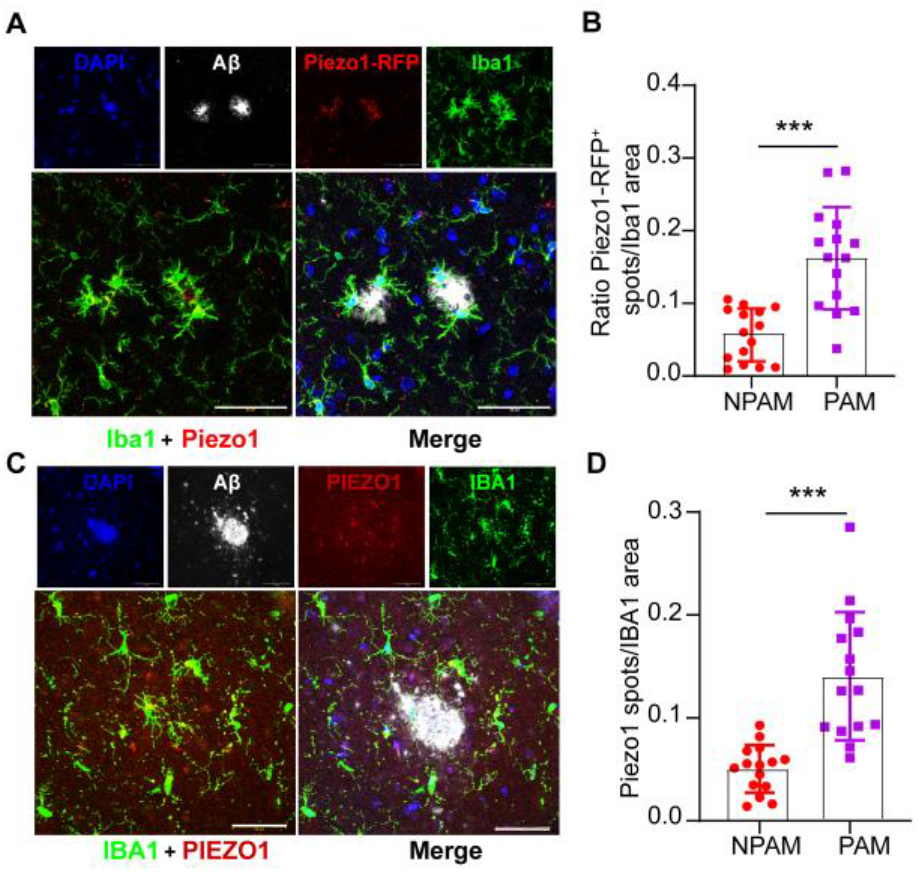
Microglial Piezo1 Expression in 5×FAD mice and AD Brain Tissue. (A) Representative images of Piezo1 (red), microglia (Iba1, green), and Aβ plaques (anti-MOAB2, white) in 5-month Piezo1^P1-tdT^;5×FAD mice. Scale bar, 50 μm. (B) Quantification of the Piezo1-RFP fluorescence spots/Iba1 area in cortical microglia of the Piezo1^P1-tdT^;5×FAD mice reveals marked Piezo1 upregulation in PAM, compared with NPAM (n = 15 independent view fields per group). (C) Representative images of Piezo1(red), IBA-1(green), and Aβ plaques (anti-MOAB2, white) in the cortex of the AD patient. Scale bar, 50 μm. (D) Each point represents the Piezo1 fluorescence spots / Iba1 area from some single microglia in PAM or NPAM in AD patients (n = 15 independent view fields per group). Statistics: Mean ± SEM. unpaired Student’s t-test (B and D); ns, no significance; ***p < 0.005.

### Microglial *Piezo1* knockout exacerbates AD-like pathologies in 5×FAD mice

The above results led us to investigate whether Piezo1 is detrimental or beneficial for microglia effector functions in Aβ pathology *in vivo*. We crossed tamoxifen (TAM)-induced microglial *Piezo1* specific knockout (*Piezo1*^*fl/fl*^; *Cx3cr1*^*CreER-EYFP*^) mice with 5×FAD mice. We injected *Piezo1*^*fl/fl*^;5×FAD (*Piezo1*^*fl/fl*^; FAD) and *Piezo1*^*fl/fl*^; *Cx3cr1*^*CreER-EYFP*^;5×FAD mice (*Piezo1*^*MGKO*^; FAD) with tamoxifen (TAM) at six weeks, and AD-like pathologies were subsequently monitored (Figure 3A). TAM induction led to the significant deletion of *Piezo1* in microglia in these mice as detected by immunofluorescence staining at 18 weeks post-induction (Figure S3A). Immunofluorescence staining using MOAB2 antibody showed significantly increased Aβ plaque accumulation in hippocampal and cortical regions of *Piezo1*^*MGKO*^; FAD mice compared to control mice (*Piezo1*^*fl/fl*^; FAD) (Figures 3B-3C, Figures S3B-S3D). Microglial *Piezo1* deficiency also increased soluble and insoluble, guanidine-extracted Aβ42 in the hippocampus and cortex of *Piezo1*^*MGKO*^; FAD mice in accord with increased Aβ plaque accumulation (Figures 3D-3E). Dystrophic neuronal membranes usually surround amyloid plaques, reflecting dystrophic neurites (Condello et al., 2011). Consistent with more Aβ plaque deposition, *Piezo1* knockout noticeably increased the total area of dystrophic neurites, as detected by immunostaining with Lamp1(Figures S4A-S4B).

**Figure 3.**
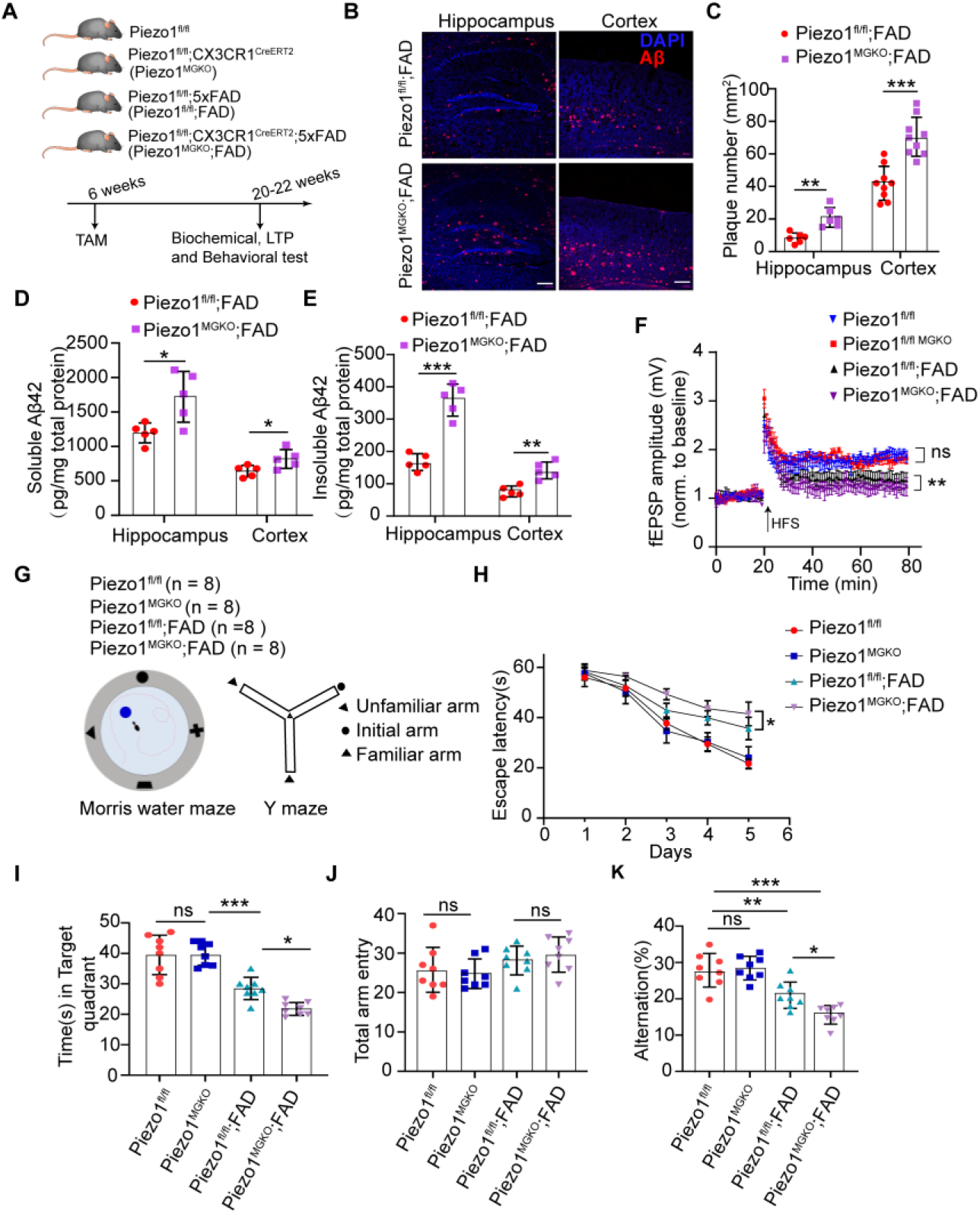
Microglial *Piezo1* Deficiency Exacerbates Aβ Pathology and Cognitive Impairment in 5×FAD mice. (A) Scheme of the experimental timeline for analyzing *Piezo1*^*fl/fl*^, *Piezo1*^*MGKO*^, *Piezo1*^*fl/fl*^; FAD, and *Piezo1*^*MGKO*^; FAD mice. (B) Representative images for Aβ (anti-MOAB2, red) in the hippocampus and cortex of 5-month-old 5**×**FAD mice indicated two groups. Scale bar, 100 μm. (C) Quantification analysis of the number of Aβ plaques in hippocampus and cortex between *Piezo1*^*fl/fl*^; FAD and *Piezo1*^*MGKO*^; FAD mice (n = 6 mice in hippocampus and n = 9 mice in cortex). (D and E) Quantitative analysis of the relative content level of soluble and insoluble Aβ42 in hippocampus and cortex measured by ELISA between the indicated groups (n = 5 mice per group). (F) LTP recording for continuous 60 min in the hippocampal Schaffer collateral region. Averaged potentiation (mean ± SEM) of baseline normalized fEPSP in the indicated groups was calculated (n = 4 mice per group). (G)Experimental scheme of adult 5**×**FAD mice and non-FAD mice for behavioral test about Morris water maze (MWM) and Y maze. (H) The escape latency time (s) of mice in each group to reach the hidden platform in the MWM test across five training days (n = 8 mice per group). (I) MWM probe test trial was performed 24 h after the training course finished, and the total time(s) in the target quadrant was recorded in each group at 5.5 months old (n = 8 mice per group). (J and K) The cognitive function of indicated groups was evaluated by the Y-maze test for measuring total arm entries (J) and spontaneous arm alternation (K) in the indicated genotypes at 5.5 months old (n = 8 mice per group). Statistics: Mean ± SEM. unpaired Student’s t-test (C, D, and E); two-way ANOVA with Bonferonni post hoc analyses (F and H); one-way ANOVA with Bonferroni post hoc test (I, J, and K); ns, no significance; *p < 0.05; **p < 0.01; ***p < 0.005.

Accumulating evidence suggests that synaptic dysfunction is the primary pathological correlate of cognitive decline in Alzheimer’s disease (Forner et al., 2017). To determine whether microglial *Piezo1* deficiency impairs synaptic function, we measured hippocampal long-term potentiation (LTP) and found that microglial *Piezo1* knockout had negligible effects on LTP in non-FAD background mice (*Piezo1*^*fl/fl*^ mice versus *Piezo1*^*MGKO*^ mice). However, in contrast to *Piezo1*^*fl/fl*^; FAD mice, there was noticeable LTP impairment in *Piezo1*^*MGKO*^; FAD mice (Figure 3F). Synapse loss correlates strongly with abnormal synaptic function and toxicity of over-deposited Aβ (Spires-Jones and Hyman, 2014). We found fewer synapses in the hippocampus of *Piezo1*^*MGKO*^; FAD mice than *Piezo1*^*fl/fl*^; FAD mice, as indicated by a decrease in the number of PSD95 (Figures S4C-S4D) and the colocalized puncta of PSD95 and synapsin (Figures S4C-S4E). No difference in synapse loss existed in non-AD mice with microglial *Piezo1* deletion (Figures S4C-S4E).

We next performed the Morris water maze (MWM) and the Y-maze tests to examine the effects of microglial *Piezo1* deficiency on spatial learning and memory in experimental mice (Figure 3G). Compared with *Piezo1*^*fl/fl*^; FAD mice, *Piezo1*^*MGKO*^; FAD mice manifest impaired learning and spatial memory in the MWM test with more time spent reaching the target platform (Figure 3H) and less preference to the target quarter at 5.5- months old (Figure 3I). Afterward, we employed the Y-maze test to acquaint working short-term memory. Consistent with their performance in the MWM, *Piezo1*^*MGKO*^; FAD had a reduced spontaneous alternation without differences in total arm entries (Figures 3J-3K). At the same time, microglial *Piezo1*-deficient mice with non-FAD backgrounds didn’t involve pervasive cognition impairment supported by MWM and Y-maze outcomes (Figures 3I and 3K). Taken together, these analyses illustrate the detrimental roles of *Piezo1* depletion in microglia in an AD mouse model with exacerbation of Aβ plaque burden, synaptic failure, and cognitive deficits.

### Microglial *Piezo1* knockout impairs the phagocytic activity of microglia

Aβ phagocytic activity of microglia directly impacts Aβ clearance both in vitro and *in vivo* (Koenigsknecht and Landreth, 2004; Pluvinage et al., 2019). Recent studies reveal that Piezo1 plays a critical role in regulating macrophage phagocytosis, enveloping and internalizing pathogens once overactivation (Geng et al., 2021; Ma et al., 2021). To investigate whether *Piezo1* deficiency in microglia could disturb cellular phagocytosis, we administered MX04 intraperitoneally to label Aβ plaques *in vivo*, and then performed immunofluorescence colocalization analysis for Iba1^+^ microglia and MX04^+^ plaques in FAD mice. We found significantly reduced Aβ engulfment by microglia in *Piezo1*^*MGKO*^; FAD mice compared to *Piezo1*^*fl/fl*^; FAD mice, as quantitative assessment of the colocalized volume of MX04 and Iba1(Figures 4A-4B). Likewise, a flow cytometry-based method to investigate microglial Aβ phagocytosis *in vivo* revealed a striking decrease in the proportion of MX04^+^ CD11b^+^ CD45^+^ microglial cells in *Piezo1*^*MGKO*^; FAD mice (Figures 4C-4D). This effect was not due to the changed numbers of microglial cells, as shown by no significant difference of CD11b^+^ CD45^+^ microglia in percentage between indicated groups (Figure S5A). These data further demonstrate that Piezo1 deficiency in microglia decreases microglial phagocytosis of Aβ *in vivo*. In addition, we found that the microglial absence of Piezo1 failed to affect the levels of amyloid precursor protein (APP), α-CTF, or β-CTF in the cortex and hippocampus of 5×FAD mice (Figures S5B-S5E), suggesting that microglial Piezo1 knockout did not influence APP metabolism.

**Figure 4.**
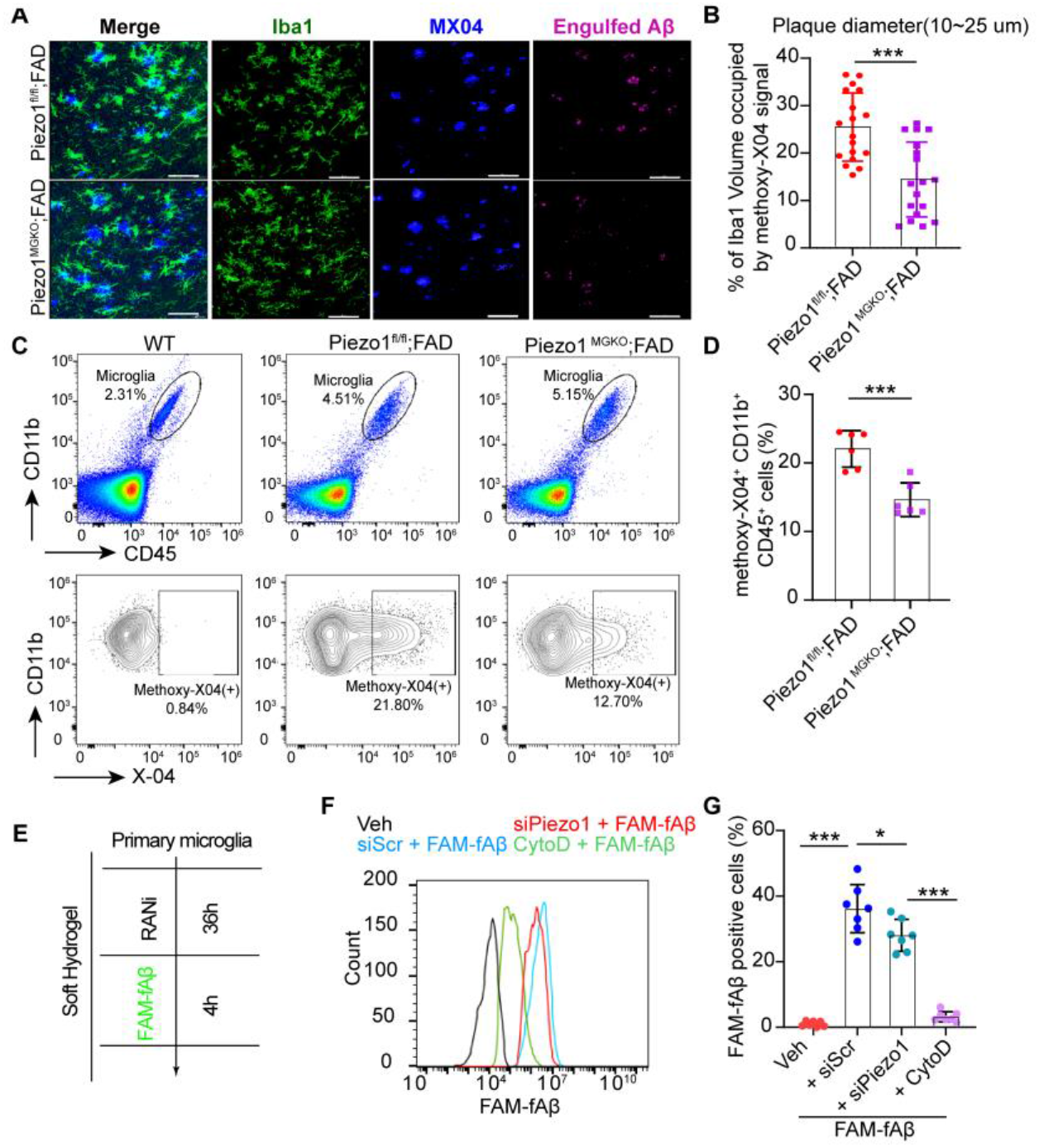
Microglial *Piezo1* Deficiency Decreases Aβ Phagocytosis. (A) Imaris based 3D reconstruction of microglia (Iba1, green) and methoxy-X04 positive (MX04^+^, blue) amyloid plaques from the hippocampus of *Piezo1*^*fl/fl*^; FAD and *Piezo1*^*MGKO*^; FAD mice at 5 months. Scale bar, 50 μm. (B) Quantification of volume of Iba1 channel colocalized with MX04 in the reconstructed images (n = 18 independent view fields per group). (C) Representative FACS dot plots showed the CD45^+^ CD11b^+^ microglia and the percentage (%) of MX04^+^ microglia by flow cytometry in the hippocampus of two indicated groups. (D) Quantification of MX04^+^ CD11b^+^ CD45^+^ hippocampal microglial cells between two groups (n = 6 mice per group). (E) Diagram was showing primary microglia phagocytosis assays. Primary microglia were seeded on soft hydrogel, accompanied by RNA interference (RNAi) for 36 h. 500 nM fibrillated FAM-fAβ was subsequently added for further FACS analysis. (F) Flow cytometry histograms of FAM-fAβ positive primary microglia treated with vehicle, si*Scr*, si*Piezo1*, and Cytochalasin D (CytoD) (10 μM), respectively. (G) Statical analysis of the percentage of FAM-fAβ positive cells between different groups to indicate the ability of FAM-fAβ42 uptake. Statistics: Mean ± SEM. unpaired Student’s t-test (B and D); one-way ANOVA with Bonferroni post hoc test (G); *p < 0.05; ***p < 0.005.

We had seen that insoluble Aβ42 fibrils as mechanical stimulus increased the Piezo1 protein level in primary microglia cultured on the soft surface. We thus suspected that Piezo1 favors microglial phagocytosis of fAβ42 once the mechanical gain of initial and enough stimulation. We determined the effects of *Piezo1* knock-down on Aβ42 fibrils phagocytic uptake in microglia seeded on soft substrate. Postnatal wild-type primary microglia cells cultured on the soft hydrogel were transfected with the *siPiezo1*, followed by FAM-fAβ42 treatment before FACS analysis (Figure 4E). We found that *Piezo1* knock-down significantly decreased FAM-fAβ42 uptake in primary microglia cells (Figures 4F-4G). In addition, fAβ42 induced cellular surface enlargement with the formation of filopodia-like structures that have been proven to facilitate phagocytosis (He et al., 2022; Kress et al., 2007) was dramatically abrogated due to *Piezo1* knock-down in primary microglia cultured on the soft hydrogel (Figures S5F-S5G). Therefore, these data illustrate that Piezo1 plays an essential role in modulating microglial phagocytic activity via stiffness sensing, which is able to contribute to the augmentation of Aβ deposition caused by microglial *Piezo1* deficiency in the brain of 5×FAD mice.

### Microglial *Piezo1* Deficiency Diminishes the Capacity of Microglia Cluster to and Organize Aβ Plaques

To further investigate the molecular mechanisms underlying the functions of Piezo1 in microglia react to Aβ plaques, we performed RNA sequencing (RNA-seq) analysis on freshly isolated primary microglia from five-month-old *Piezo1*^*MGKO*^; FAD and *Piezo1*^*fl/fl*^; FAD mice (Figures 5A and S6A). We identified 466 differentially expressed genes, with 410 downregulation and 56 upregulation in *Piezo1*^*MGKO*^; FAD versus *Piezo1*^*fl/fl*^;FAD (log_2_^|FC|^ > 1.0, p < 0.05). Several innate immune response genes were up-regulated, such as *Acod1*, *Mefv*, and *Pglyrp2*, while many cytoskeleton dynamics related genes (*Actg2*, *Acta2*, *Cttn*, *Myl9*, *Myh14*, *Cttnbp2*, etc.) were down-regulated (Figure 5B). Gene Ontology (GO) analysis of the significantly down-regulated genes in *Piezo1* conditional knockout microglia revealed a significant functional reduction in synapse organization, cell-substrate adhesion, and cytoskeleton system-based events (morphogenesis, migration, and endocytosis) in 5×FAD mice (Figure 5C), suggesting the inhibition of microglial adhesion signaling and cytoskeleton-associated signaling in *Piezo1* knockout microglia. On the other side, no significantly enriched GO terms were found among the up-regulated genes. In addition, we performed gene set enrichment analyses (GSEAs) and found that microtubule-based movement and substrate-dependent cell migration signaling were significantly down-regulated in microglia in *Piezo1*^*MGKO*^; FAD mice (Figure 5D). Furthermore, Ingenuity Pathway Analysis (IPA) revealed that microglial *Piezo1* deficiency was associated with decreased pathways of calcium signaling, actin cytoskeleton-based signaling, and signaling by Rho family GTPase (Figure S6B) that are important for mechanotransduction (Nourse and Pathak, 2017; Ohashi et al., 2017; Orr et al., 2006). In sum, these data elucidate that in the presence of Aβ plaques, *Piezo1* deficiency mainly derails the Piezo1-actin cytoskeleton pathway in microglia, an important pathway to transduce mechanical stimulations (Nourse and Pathak, 2017), which probably impairs microglial responses to Aβ plaque stiffness.

**Figure 5.**
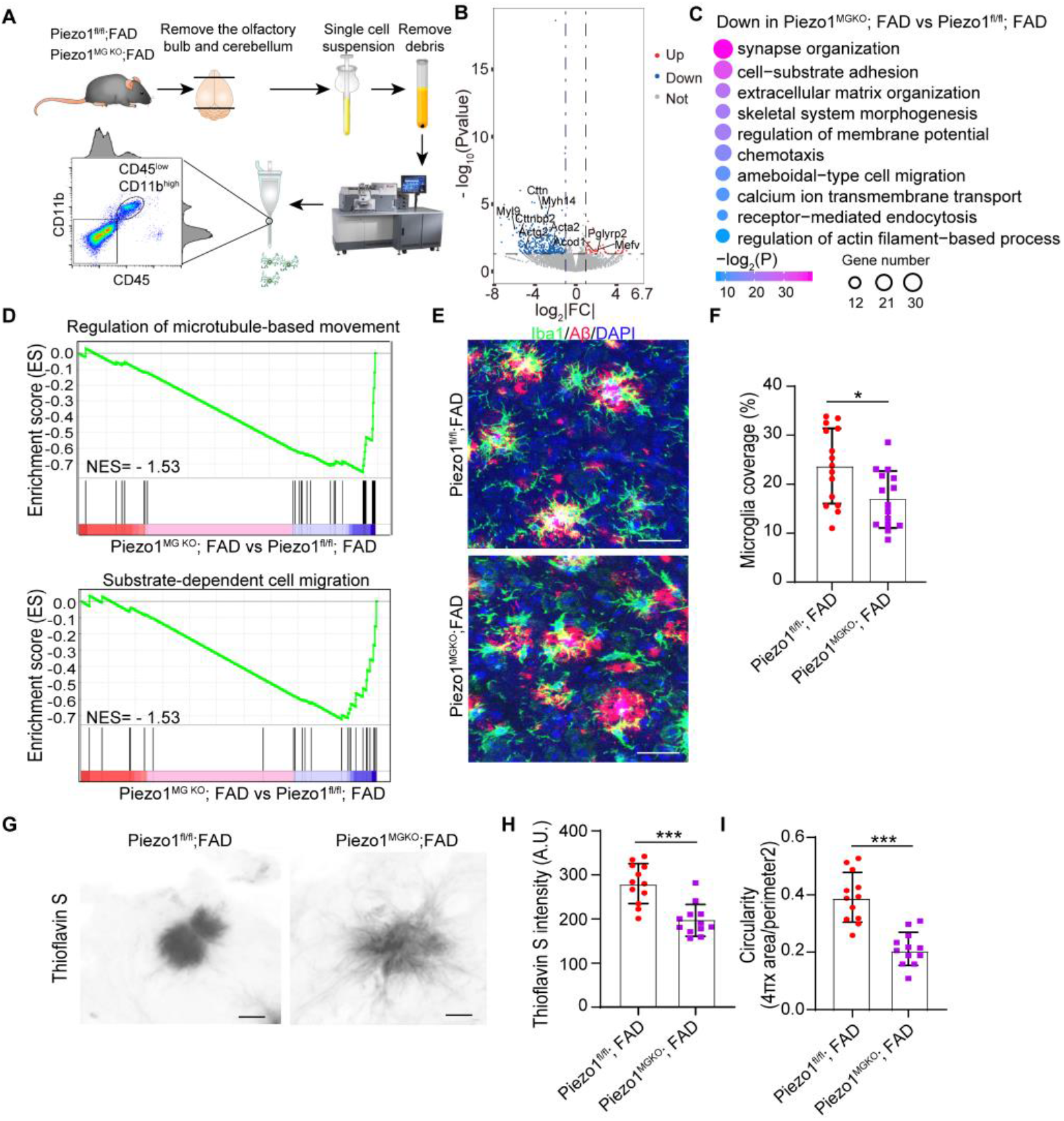
*Piezo1* Deficiency Programs Microglia to Reduce Clustering of Aβ Plaques and Plaque compaction. (A) Schematic of the methodology for isolating CD45^low^ CD11b^high^ microglia from the indicated groups at 5-month-old to perform RNA-seq. (B) Volcano plot illustrating differential expression of genes in microglia isolated from *Piezo1*^*MGKO*^; FAD versus *Piezo1*^*fl/fl*^; FAD. (C) GO enrichment analysis of downregulated genes in *Piezo1*^*MGKO*^; FAD versus *Piezo1*^*fl/fl*^; FAD. (D) GSEA analysis showing the enrichment score of regulation of microtubule-based movement and substrate-dependent cell migration pathways in microglia between Piezo1 ^MGKO^; FAD versus Piezo1^fl/fl^; FAD. (E) Representative confocal images show the interaction between Aβ plaques Aβ (anti-MOAB2, red) and microglia (Iba1, green) at 5-month-old indicated genotypes. (F) Quantification of microglia coverage (%) between two groups (n = 15 independent view fields per group). (G) Morphology of Thioflavin S reactive Aβ plaques was visualized by high-resolution confocal microscopy. Scale bar, 5 μm. (H and I) Qualification analysis of Thioflavin S intensity in (H) and circularity in (I) (n = 12 independent view fields per group). Statistics: Mean ± SEM. unpaired Student’s t-test (F, H, and I); *p < 0.05; ***p < 0.005.

Given these findings, we then characterize the role of Piezo1 in shaping microglial motility and distribution in 5×FAD mice. We found that *Piezo1* knockout reduced microglial clustering of Aβ plaques *in vivo*, as shown by the decrease in the microglial volume coverage surrounding Aβ plaques (Figures 5E-5F). In addition, Aβ plaques displayed an irregular shape with more fiber-like structures extending outwards and lacked the homogenous staining intensity in microglial *Piezo1* deficiency 5×FAD mice. As such, both mean Thio-S intensities and plaque circularity were reduced in 5×FAD mice devoid of microglial *Piezo1*(Figures 5G-5I). Overall, these data indicate that the Piezo1-mediated microglial mechanical responses are crucial in segregating the plaques, preventing plaque expansion, and facilitating its compaction.

### Piezo1 activation ameliorates AD-like pathologies in 5×FAD mice

Since microglial *Piezo1* deficiency leads to worsening Aβ burden and cognitive impairment in 5×FAD mice, we thus investigated whether activation of Piezo1 altered these AD-like pathologies in 5×FAD mice. Firstly, we sought to determine the pharmacokinetic properties of the Piezo1 specific agonist Yoda1 in mice (Figure S7A). We found that Yoda1 reached a maximum concentration in the brain and plasma at 0.5 hours after a single administration of Yoda1(3.0 mg/kg) by intraperitoneal (i.p.) injection, which was maintained for up to 15 hours (Figures S7B-S7C). We then administered Yoda1 to 1.5-month-old 5×FAD mice (Figure 6A). Mice treated for 3.5 months with Yoda1 did not exhibit liver or bodyweight change gain compared to vehicle (corn oil) control (Figures S7D-S7F), indicating no apparent toxic side effects of Yoda1. Notably, 5×FAD mice administrated Yoda1 had less Aβ plaque burden (Figures 6B-6D), in line with fewer brain Thio-S^+^ Aβ plaques (Figures S7G-S7H), and lowered hippocampal soluble and insoluble Aβ42 levels in 5×FAD mice after Yoda1 administration (Figures S7I-S7J). Besides, both Lamp1^+^ dystrophic neurites (Figures 6B and 6E) and SMI31^+^ neurofilament-containing dystrophic neurites in the brain of 5×FAD mice (Figures S7K-S7L) diminished significantly with Yoda1 treatment. Interestingly, we found that Yoda1 treatment caused more recruitment of microglia to Thio-S^+^ Aβ plaques in 5×FAD mice than vehicle (corn oil) treatment (Figures 6F-6G). Consequently, the amount of Aβ internalized by microglia was significantly more tremendous in Yoda1-treatment mice than in vehicle-treatment mice (Figures 6H-6I). To investigate whether reduced Aβ accumulation in 5×FAD mice after Yoda1 administration led to cognitive improvement, we also performed the MWM test. In the acquisition phase, 5×FAD mice showed significant learning disability compared to non-transgenic wild-type (WT) mice, while this phenotype was partially ameliorated in 5×FAD mice after Yoda1 administration (Figures 6J-6K). As assessed in probe tests, administration of Yoda1 in 5×FAD mice spent more time in the target quadrant compared to control 5×FAD mice, indicating a better memory function in 5×FAD mice after Yoda1 administration (Figure 6L). Collectively, these results demonstrate that chronic Piezo1 activation by Yoda1 administration of 1.5-month-old 5×FAD mice for 3.5 months ameliorates Aβ deposition and behavioral deficits likely via modulation of the microglial function to some extent in 5×FAD mice.

**Figure 6.**
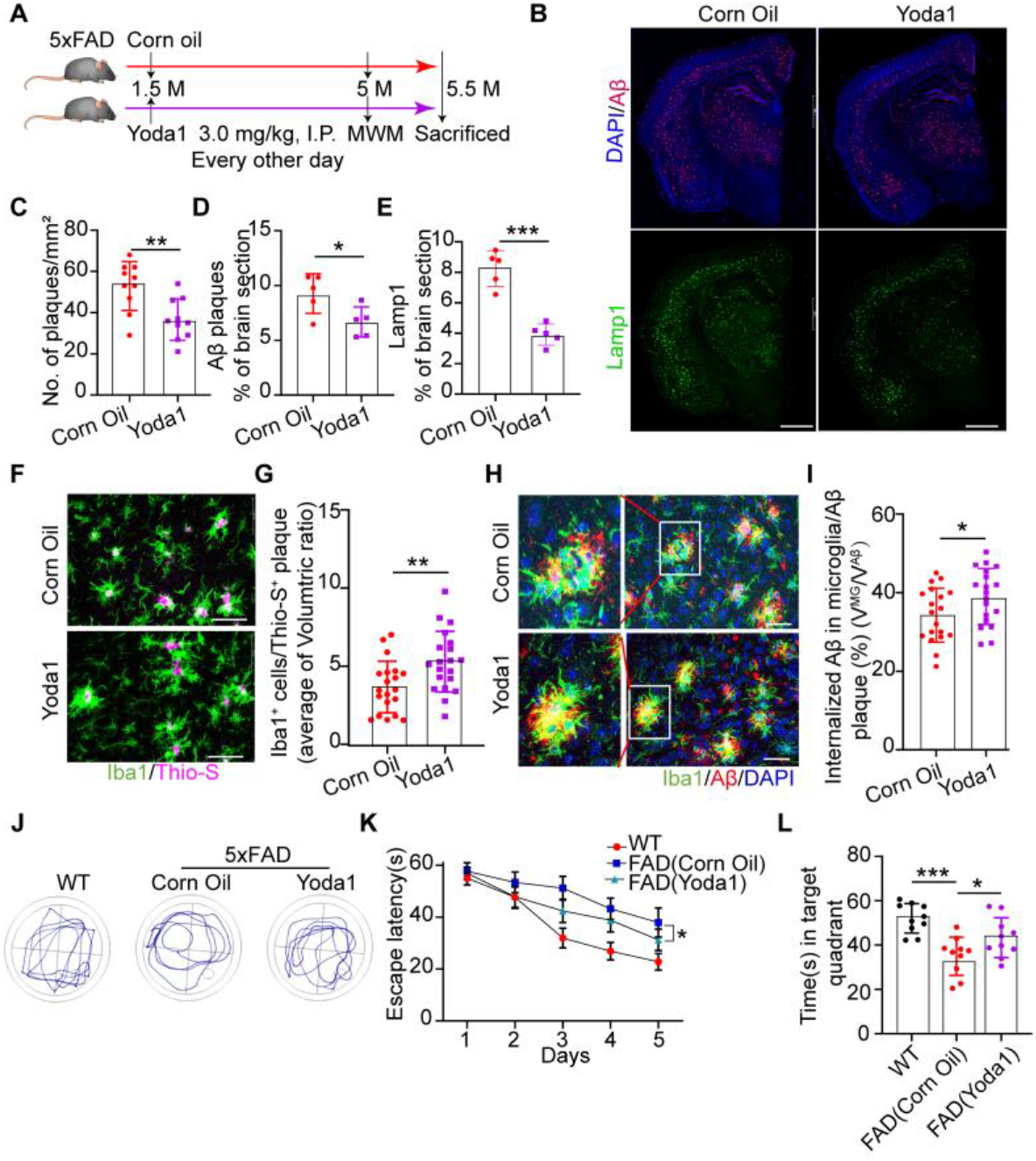
Yoda1 Administration Ameliorates the Aβ Burden and Cognitive Decline in 5×FAD mice. (A) Timeline of Yoda1 injection experiments. 5×FAD mice were injected intraperitoneally with Yoda1(3 mg/kg) or vehicle (Corn oil) every other day at the 1.5-month-old. (B) Representative images of brain sections from 5xFAD mice treated with Yoda1 or oil for staining of Aβ (anti-MOAB2, red) and LAMP1(green). Scale bar, 1 mm. (C) The Number of Aβ plaques per mm^2^ (n = 10 independent view fields per group). (D and E) Aβ plaques and Lamp1 coverage in (B) (n = 5 mice per group). (F) Representative images of brain stained for dense-core plaques (Thio-S, purple) and microglia (Iba1, green). Scale bar, 50 μm. (G) The average volumetric ratio per Aβ plaque was qualified (n = 20 plaques from three independent experiments). (H) Aβ and microglia are indicated by immunostaining for the antibodies MOAB2 and Iba-1, respectively. Scale bar, 50 μm. (I) Internalized Aβ in microglia/Aβ plaque (%) was calculated. V^MG^ (total volume of plaque-associated microglia) and V^Aβ^ (the volume of Aβ plaques) (n = 19 independent view fields per group). (J and K) Morris water maze tests were performed in the invisible platform training (n = 10 mice per group). (L) MWM probe test results about time(s) in target quadrant (n = 10 mice per group). Statistics: Mean ± SEM. unpaired Student’s t-test (C, D, E, G, and I); two-way ANOVA with Bonferonni post hoc analyses(K); one-way ANOVA with Bonferroni post hoc test (L); *p < 0.05; ***p < 0.005.

## Discussion

Given the multiple chemical composition of the Aβ plaque, previous work has focused on the chemical property of the Aβ plaque that drives the microglia response (Drummond et al., 2017; Stewart and Radford, 2017). Physical measurements in the formalin-fixed AD mouse brain slices reveal that Aβ plaque has a rigid core formed by insoluble Aβ fibrils (Mattana et al., 2017). Microglia, the brain-resident immune cells, mount immune responses to various brain injuries, including Aβ deposition in AD (Hansen et al., 2018; Madore et al., 2020). Recently studies indicate that microglia can respond to changes in mechanical forces and preferentially migrate towards stiffer substrates (Bollmann et al., 2015; Meller et al., 2017), whereas mechanosensation in microglia reaction to Aβ plaque stiffness is unclear. In this project, we highlight the involvement of the mechanically gated ion channel Pizeo1 in Aβ fibrils stiffness sensing by microglia and their responses to brain Aβ accumulation and cognitive function in an AD mouse model.

Although Aβ fibrils exhibit high stiffness, the actual mechanical feature of the Aβ plaque-associated tissues (PAT) *in vivo* is still unexplored. Combined AFM-fluorescence microscope and MX04 labeling Aβ plaques *in vivo*, we discover the PAT’s stiffness (582.8 ± 109.9 Pa) is not tremendous but still significantly higher than the non-Aβ plaque-associated tissues (NPAT, 317.9 ± 33.78 Pa) in acute horizontal live 5×FAD mouse brain slices. Intriguingly, AD brain stiffness tends to decrease on the macroscale, likely due to widespread neuronal cell loss (Hiscox et al., 2020). It is plausible that the disparity between the stiffness features of the PAT and NPAT may evoke microglial mechanosensation of the Aβ plaque in AD progression. Evidence shows that Piezo1 mediates varied environmental stiffness sensing in multiple cell types, including myeloid cells, neural stem cells, and oligodendrocyte progenitor cells (OPCs) (Pathak et al., 2014; Segel et al., 2019; Solis et al., 2019). Noticeably, as we and others report, Piezo1 is the highest expressed in microglia among the previously reported mechanosensory ion channels (Galatro et al., 2017; Zhang et al., 2014). However, it has not been clear whether Piezo1 involve in microglial mechanosensation. Our data show that primary microglia seeded on a stiffer polyacrylamide hydrogel elicits a significant elevation of the Piezo1 protein level. Intriguingly, the Piezo1 protein level increases upon fAβ42 stiffness stimulation instead of its chemistry cue, and fAβ42 stiffness-dependent increases in Ca^2+^ influx activity required Piezo1 in primary microglia.

Furthermore, we find a distinctly higher level of the Piezo1 protein in microglia around Aβ plaque than those away from plaque both in the brain of 5×FAD mice and AD patients, demonstrating that Piezo1 may also prime microglial sensing of the Aβ plaque stiffness *in vivo*. Although the exact mechanisms that activate Piezo1 in response to insoluble Aβ stiffness warrant further studies, our results suggest that Piezo1 links the microglial mechanosensation of the insoluble Aβ aggregates *in vitro* and *in vivo*. Next, we discern the function of the Piezo1 in microglia that involves in Aβ pathology. We confirm that microglial *Piezo1* knockout dramatically exacerbates brain Aβ plaque burden and cognitive impairment in 5×FAD mice but not in non-FAD subjects, suggesting Piezo1-dependent microglial mechanosensation may protect against Aβ pathology.

Previous work has concentrated on characterizing receptor-mediated chemical signaling pathways in microglial phagocytosis, but mechanosensation in microglial phagocytosis is unclear. Mechanosensation by Piezo1 is essential for innate immune functions, such as phagocytosis (Aykut et al., 2020; Geng et al., 2021; Ma et al., 2021). Our finding is that microglial *Piezo1* deficiency dramatically reduces Aβ plaque material phagocytosis by microglia in the brain of 5×FAD mice. Subsequently, in vitro data show that siRNA-mediated *Piezo1* knock-down impairs fAβ42 uptake in primary microglia seeded on soft substrate. Microglia engulfing Aβ fibrils requires cytoskeletal rearrangements (Koenigsknecht and Landreth, 2004). Indeed, *Piezo1* knock-down disturbs cytoskeletal reorganization following fAβ42 stimulation in primary microglia. Furthermore, RNA sequencing of microglia freshly isolated from *Piezo1*^*fl/fl*^; FAD and *Piezo1*^*MGKO*^; FAD mice brains identifies significant down-regulated genes responsible for cytoskeleton system-based signaling in microglia from *Piezo1*^*MGKO*^; FAD mice. A recent study indicated that microglial polarization responses to stiff tissue require Kindlin3 (Dudiki et al., 2020), an intracellular adaptor protein, recruiting cytoskeletal proteins involved in cell adhesion and migration (Zhao et al., 2012). Piezo1 channel activity triggers many intracellular processes involving the actin cytoskeleton (Nourse and Pathak, 2017). Hence, whether Piezo1 may be an upstream regulator of Kindlin3 warrants further investigation. In addition, Piezo1 as a sensor for mechanical stimuli regulates downstream calcium signaling during phagocytosis (Ma et al., 2021), which is predominantly downregulated in *Piezo1*-deficient microglia in 5×FAD mice as IPA analysis displays. Therefore, Piezo1-mediated mechanotransduction is indispensable for microglial phagocytosis of Aβ plaques materials.

Microglia surrounding the Aβ plaque surface constitutes a barrier that limits the expansion of Aβ plaque, facilitates its compaction, and ameliorates plaque-induced neurotoxicity (Condello et al., 2015; Yuan et al., 2016; Zhao et al., 2017).In *Piezo1*^*MGKO*^; FAD mice, we identify that *Piezo1* deficiency impairs the ability of microglia to cluster around the Aβ plaques and leads to more diffuse plaques. It is possible that deletion of the microglial *Piezo1* disrupts its durotaxis, which explains the reduced recruitment of microglia to the Aβ plaques. In addition, GO enrichment analysis shows that the downregulated genes in microglia isolated from *Piezo1*^*MGKO*^; FAD mice are mainly enriched in cell adhesion. Cell adhesion is relevant for structural integrity and cell barrier formation (Sumigray and Lechler, 2015). Therefore, this impaired microglial adhesion function may reduce the microglia barrier build-up around Aβ plaques and promote microglial Aβ propagation in *Piezo1*^*MGKO*^; FAD mice as a new study confirms that microglia contribute to Aβ propagation into unaffected brain regions *in vivo* (d’Errico et al., 2022). In summary, Piezo1-mediated mechanical responses are required for microglial phagocytosis and compaction of the Aβ plaque, limiting Aβ pathology in the brain.

Yoda1 is a small synthetic molecule that can selectively activate the Piezo1 channel without mechanical stimuli (Syeda et al., 2015). We confirm that Yoda1 is BBB permeability and long-term systemic administration of Yoda1 decreases brain Aβ deposition and plaque-associated neuritic dystrophy, ultimately improving cognitive function in 5×FAD mice. Notably, Yoda1 treatment causes more microglia clustering around Aβ plaques and higher amounts of Aβ internalized by microglia than vehicle treatment in 5×FAD mice. These data display that Yoda1 induced Piezo1 channel activation reverses the Aβ pathology *in vivo*, probably by enhancing Piezo1-mediated microglial functions. However, we cannot rule out the possibility that such an approach would be the functions of Piezo1 in other tissues or other brain cells. For example, astrocytes are generally devoid of Piezo1, while exhibiting up-regulated Piezo1 expression in astrocytes around Aβ plaques in postmortem AD brains and a rat model of AD (Satoh et al., 2006; Velasco-Estevez et al., 2018). Future studies will be needed to clarify whether Piezo1 in these non-microglia cells is functionally linked to Aβ pathology *in vivo*.

In conclusion, our study indicates that insoluble Aβ aggregates present a mechanical stimulus to microglia, which prevents the spreading of Aβ pathology through mechanosensation by microglial Piezo1. Moreover, the chemical activation of Piezo1 mitigates AD-like pathology, raising the probability that modulating Piezo1 activity might have therapeutic value for AD. Further exploring the exact mechanotransduction pathway via microglial Piezo1 will expand our comprehension of how microglia contribute to the pathogenesis of AD.

## Methods

### Human brain tissue

The Human brain tissue was provided by National Health and Disease Human Brain Tissue Resource Center. The detailed information of human brain samples of AD was represented in Figure S2C.

### Animals

All animal experiments were approved by Institutional Animal Care and Use Committed at Xiamen University. Mice were housed in a 12:12-h light-dark cycle with ad libitum access to food and water. *Piezo1*^*fl/fl*^ mice (Stock no. 029213) and *Piezo1*^*P1-tdT*^ mice (Stock no. 029214) were kindly provided by Dr. Dawang Zhou (Geng et al., 2021) at Xiamen University. Wild-type C57BL/6N mice, *Cx3cr1*^*CreER-EYFP*^ mice (Stock no. 021160) and 5×FAD mice (MMRRC Strain no. 034840-JAX) were initially from The Jackson Laboratory. *Cx3cr1*^*CreER-EYFP*^ mice were crossed with *Piezo1*^*fl/fl*^ mice to generate *Cx3cr1*^*CreER-EYFP*^; *Piezo1*^*fl/fl*^ mice. Then, 5×FAD mice were crossed with *Cx3cr1*^*CreER-EYFP*^; *Piezo1*^*fl/fl*^ mice to develop *Cx3cr1*^*CreER-EYFP*^; *Piezo1*^*fl/fl*^; 5×FAD. *Cx3cr1*^*CreER-EYFP*^; *Piezo1*^*fl/fl*^; 5×FAD and *Piezo1*^*fl/fl*^; 5×FAD male littermates aged at 4.5 weeks were subjected with four daily intraperitoneal 20 mg/kg tamoxifen (TAM, dissolved in corn oil (Solarbio), Beidapharma Co., Ltd) to generate microglial *Piezo1* knockout in 5×FAD mice (*Piezo1*^*MGKO*^; FAD) and the control littermate mice (*Piezo1*^*fl/fl*^; FAD).

### Polyacrylamide hydrogel preparation

The details of polyacrylamide gels of varying stiffness refer to Segel et al.’s research (Segel et al., 2019). Circle microscope coverslips (NEST) were divided into bottom coverslips and top coverslips. The bottom coverslips were coated with 1.2 % bind silane (Sigma) dissolved in 95% ethanol and 5% glacial acetic acid and air-dried for further use. The top coverslips were incubated in 15% Sigmacote (Sigma) dissolved in chloroform (Sigma) for 1 h. Then, the top coverslips were air-dried and polished cleanly. To make the soft gel, 7 % acrylamide (Sigma), 8 % bis-acrylamide (Sigma) and 20 % 6-acrylamidohexanoic acid (Aladdin) in ddH_2_O with 0.004 g ml^−1^ TEMED (Sigma) and 0.001 g ml^−1^ ammonium persulfate (Sigma). To make the stiff gel, acrylamide was mixed with bis-acrylamide equally, followed by adding TEMED and ammonium persulfate. The mixed acrylamide solution was rapidly pipetted onto the bottom coverslip, and the top coverslip was placed on the semi-solidified hydrogel. Then the top coverslip was removed, and polymerized hydrogel adhered to the bottom coverslip.

To activate the hydrogels, hydrogels were first washed twice with methanol, rehydrated in PBS, and incubated in MES working buffer (4-morpholineethansulfonic acid (MES) hydrate (Sigma), 500 mM NaCl (Sigma), pH = 6.1) for 10 min. Next, hydrogels were activated with activate buffer (480 mM N-hydroxysuccinimide (Sigma), 200 mM N-(3-dimethylaminopropyl)-N′-ethylcarbodiimide hydrochloride (Sigma) dissolved in MES working buffer). Activated hydrogels were incubated in 50 μg ml^−1^ laminin (Sigma) and Fibronectin (Novoprotein, 1μg/ cm^2^) diluted in 1×poly-L-lysine buffer overnight. Coated hydrogels were incubated and gently repeated-washed in DMEM for 3 days and exposed in UV irradiation for 1 h before usage.

### Preparation of mAβ and fAβ

Unlabeled and FAM-labeled synthetic human Aβ_1-42_ peptides (Anaspec) were dissolved in 1,1,1,3,3,3-hexafluoro-2-propanol (HFIP) (Sigma), followed by evaporation under a gentle stream of oxygen-free nitrogen gas, and the dried peptides were stored at −80°C. For Aβ monomers (mAβ) preparation, dried Aβ1-42 was dissolved with DMSO at a concentration of 1 mM and used immediately. Aβ fibrils (fAβ) were prepared by diluting the mAβ solution to 100 μM in DMEM then incubated at 37°C for 3~5 days to fibrillation.

### Primary microglia culture

Primary microglial cultures were prepared as described previously (Atagi et al., 2015). Briefly, brains were removed from non-transgenic wild-type (WT) mice at postnatal days 2-3. After removing the meninges and cerebella, the remaining brain tissues were triturated by pipetting several times in the ice-cold HBSS buffer (Solarbio). Mixed glial cells were plated onto poly-L-lysine-coated flasks and grown in DMEM supplemented with 10% fetal bovine serum (FBS) (Gibco). After a 24 h incubation, the medium was changed to that which contained 25 ng/ml mouse GM-CSF (R&D Systems). The primary microglia were harvested by shaking (200 rpm, 30 min) 10-14 days after plating and subjected to various treatments within 24 h of harvest.

### The sample preparation for AFM

For AFM measurements of Aβ plaques-associated tissue and non-Aβ plaques-associated tissue in the brain, 5-months 5×FAD mice were pre-injected intraperitoneally with methoxy-X04 (10 mg/kg, Toris, dissolved in 10% DMSO/90 % PBS (pH 12)). Mice were anesthetized with isoflurane and quickly decapitated. The brain was dissected out and transferred into 4 °C artificial cerebrospinal fluid bubbled with 95% O_2_ and 5% CO_2_ (aCSF, containing 26 mM NaHCO_3_, 1.25 mM KH_2_PO_4_, 2 mM NaOH ,1 mM CaCl_2_, 4 mM MgSO_4_, 191 mM sucrose, 20 mM glucose, 3 mM myoinositol, 0.75 mM K-gluconate, 5 mM ethyl pyruvate, 2 mM kynurenic acid and 1 mM (+)-sodium L-ascorbate) (Koser et al., 2015). Next, the brain was fixed and cut 250-μm thick coronal brain sections with Vibrating Microtome (LEICA VT1000S) in cold-aCSF. Then, brain sections were glued at the bottom of a 35-mm glass-bottom Petri dish with Cell-Tak-coated (Corning), constantly perfused with fresh aCSF bubbled with 95% O_2_ and 5% CO_2_ in steady. The petri dish was mounted on an inverted fluorescence microscope (Zeiss) for imaging Aβ plaques with 405 nm excitation.

For AFM measurements of fAβ_1-42_ in vitro, 200 μM fAβ_1-42_ stock solution was obtained and incubated at 37 °C for 3 days. Before measurement, the fAβ_1-42_ stock solution was diluted into 20 μM with HEPES buffer and dropped onto AFM mica. Next, fAβ_1-42_ coated mica was washed to remove excessive aggregates with ddH_2_O and dried adequately for AFM detection.

### Atomic force microscopy

AFM indentation measurements were performed with an atomic force microscope (Bruker Nanowizard4). For AFM measurements of brain sections or soft hydrogel, tipless silicon cantilever (probe: NP-010; spring constant = 0.24 N m^−1^; BRUKER) was glued with polystyrene beads (diameter = 22 μm; microParticles GmbH) to the tip of the cantilever. For stiff hydrogel, a tipless silicon cantilever (probe: FESP-V2; spring constant = 2.8 N m^−1^; BRUKER) was used. Cantilever spring constants were determined by AFM software (JPK SPM). Force–distance curves were taken with the set force of 5 nN and speed of 2 μm s ^−1^ by force spectrum model for brain or soft hydrogel measurement and stiff hydrogel. To measure the fAβ_1-42_ sample, a silicon cantilever (probe: FESP-V2; spring constant = 2.8 N m^−1^; NanoWorld) was used and scanned with a QI – model with a set force of and speed of 100μm/s. The elastic moduli were extracted from force-distance curves by Hertz model (Lin et al., 2009). As previous research reported, small deviation and reliability about the Hertz model were calculated and assessed with MATLAB (Mathworks) (Zhang et al., 1997). And shear modulus of polyacrylamide gels was calculated by referring to Moshayedi et al. reported (Moshayedi et al., 2010).

### Real-time quantitative PCR analysis

Total RNA was isolated using TRIzol reagent (Ambion), and 1000 ng total RNA was reversed to cDNA using 5×All-In-One RT MasterMix (Abmgood). Quantitative PCR was performed on CFX Connect Real-Time PCR Detection System using ChamQ Universal SYBR qPCR Master mix (Vazyme) according to manufacturer’s instructions. The 2^−ΔΔCt^ method was used to calculate relative expression level after referring to β-actin as an internal control. The primer sequences were listed as follows:

Piezo1-Forward:CTTACACGGTTGCTGGTTGG;

Piezo1-Reverse:CACTTGATGAGGGCGGAAT;

Piezo2-Forward:CATGACCTCTGCCTCCATCA;

Piezo2-Reverse:CCAGGGAAAGAAGCCGAACT;

Stoml3-Forward:TCGAATCCCGACCCAGGAG;

Stoml3-Reverse:GGCTTTTGTTGGTGCCCACT;

Kcnk4-Forward:GCGATTATGTACCCGGCGATG;

Kcnk4-Reverse:CTGAGGCGAAGTAGGCTAGG;

Trpv4-Forward:GAGTCCTCAGTAGTGCCTGG;

Trpv4-Reverse:CAACAAGAAGGAGAGCAGTC;

Trpa1-Forward:GATGCCTTCAGCACCCCATTG;

Trpa1-Reverse:CACTTTGCGTAAGTACCAGAGTGG;

β-actin-Forward:GACCCAGATCATGTTTGAGA;

β-actin-Reverse: GAGCATAGCCCTCGTAGAT

### RNA interference

Small interfering RNAs (siRNAs) (Scrambled Control siRNA and Piezo1 siRNAs, Ribobio) were dissolved in 5×siRNA buffer (Dharmacon). Primary microglia were transiently transfected with siRNAs using Lipofectamine™ RNAiMAX (Thermo Fisher Scientific) with a final concentration at 40 nM following the manufacturer’s protocol. 24~48 h after siRNAs were transfected, the culture medium was replaced for further assays.

### Ca2+ imaging and analysis

Primary microglia seeded on soft was undergone corresponding stimulus interventions. After interventions finished, primary microglia of indicated groups were cultured in heat-inactivated DMEM medium containing 5 μM Fluo-4 AM (AAT Bioquest), 0.4 % F-127 (Beyotime) for 30 min at 37 °C, followed by washing twice gently with DMEM. Then, incubate cells with HBSS containing Ca^2+^ and Mg^2+^ ions (Solarbio) for 30 min at 37 °C in an incubator and 30 min at room temperature (RT) away from light for environmental adaptation, respectively. Next, the dish was real-time monitored with Leica confocal microscope (Zeiss LSM980+Airyscan2) for 5 min with a 500 ms frequency interval. Individual Ca^2+^ events were acquired with dynamic fluorescence intensity (FI) according to each time point ROI from Leica confocal microscope. The baseline FI of Ca^2+^ was defined as F°. The real-time fluorescence intensity was defined as F. Finally, [F / F°] was used to assess Ca^2+^ activity.

### Brain stereotaxic injection of FAM-fAβ_1-42_

*Piezo1*^*P1-tdT*^ mice at 4 months were anesthetized and fixed in the stereotaxic injection apparatus (RWD Life Science). After skin incision, FAM-fAβ1-42 (Anaspec, 2μg in 2μL vehicle/side) was unilaterally injected into the hippocampus (from bregma, anterior posterior: −2.2 mm; anterior posterior: ± 2 mm, dorsal ventral: −1.45 mm) using a microliter syringe (Hamilton) in 5 mins. The needle was removed after 10 mins from the completion of the infusion. Mice were sacrificed for cryostat sectioning and staining 24 h after injection.

### Immunohistochemistry, imaging, and analysis

For Immunofluorescence of the mouse brain sections, mouse brains were perfused with cold 1×PBS, followed by 4% PFA. Fixed mouse brains emerged in 20% and 30% sucrose, respectively. After the mouse brains were fully dehydrated, coronal or sagittal sections were cut on a cryostat at 20 μm thick. Sections were pre-dried for 30 min, washed with 1×PBS, and fixed in 2% PFA for 15 min at RT. Then sections were applied with antigen retrieval (citrate buffer (pH 6.0))and blocked with 3% BSA and 0.4% Triton X-100. Primary antibodies incubation overnight at 4 °C and Alexa-fluorophore conjugated secondary antibodies for 2 h at RT. For Immunofluorescence of the human brain sections. Dewaxed paraffin sections after air dry, fix slides with 2 % PFA at RT for 15 min. Block brain tissue with PBT (0.3 % Triton, 5% normal donkey serum(Yeasen) in 1×PBS) at RT for 1 h. Then brain sections were incubated with primary antibodies overnight, followed by secondary antibodies for 6 h at RT. Wash slides in PBT for 10 min, followed by 2% PFA fixation for 15 min. Stain the slides in 0.3 % Sudan Black B(Shyuanye) for 20 min. Sections were sealed with an anti-fade reagent (VECTASHIELD) for further imaging (Leica confocal microscope TCS SP8 DLS). For Thioflavin-S staining, frozen mouse brain sections were incubated with 1 mg/ml Thioflavin-S (Sigma) for 7 min. Wash with 70%, 80%, and 90% ethanol, respectively. Thioflavin S-labeled amyloid plaques’ morphology was further imaged with high-resolution confocal microscopy(Zeiss confocal microscope (LSM 880+Airyscan)) at 60× magnification with 1024×1024 pixel resolution, z-step size of 5 μm. Images were further analyzed by ImageJ software. To analyze the colocalization of Aβ plaque and microglia, 20-μm z-stacks were performed with 1.0-μm steps in the z-direction, and 1024 x 1024-pixel resolution images were analyzed by Bitplane Imaris (x64 9.2.0). The Imaris function “Surface” was used to quantify the percentage of microglia volume occupied by the Aβ plaque signal. The following antibodies of immunohistochemistry staining were used in this study: anti-Aβ (MOAB-2, Abcam, ab126649), anti-Iba1 (Wako, 019-19741), anti-Iba1 (Abcam, ab5076), anti-Piezo1 (Proteintech, 28511-1-AP), anti-Lamp1 (ABclonal, A16894), anti-RFP (Rockland, 600-401-379), anti-Smi31(Biolegend, 801602), anti-PSD95(Abcam, ab12093), anti-Synapsin(Ruiying Biological, RLT4483).

### Western blotting

Brain tissues of cortex and hippocampus were homogenized using in ice-cold RIPA buffer (150 mM NaCl,50 mM Tris-HCl, 0.1% SDS, 1% Triton X-100, 0.5% sodium deoxycholate, Phosphatase and Protease Inhibitor Cocktail (Abcam), pH 7.6) and lysed at 4°C for 30 min. After lysate was centrifuged at 12000 g at 4°C for 10 min, the supernatants containing proteins were qualified by BCA Protein Assay Kit (Vazyme) and then subjected to SDS buffer. Primary microglia seeded on various stiffness hydrogels were discarded culture medium and washed gently with cold 1× PBS. 1×SDS buffer was directly added to lysate cells adequately. The proteins were separated by 4-20% SDS-PAGE (GenScript) and transferred onto PVDF membranes(Amersham). After blocking with 5% nonfat milk in TBST (Tris-buffered saline with 0.1% Tween 20) at RT for 1 h, membranes were washed 3 times with TBST for 15 min, and incubated primary antibodies overnight. Chemiluminescent peroxidase-labeled secondary antibodies (ABclonal) were incubated for 1 h at RT. After 30 min of washing in TBST, the protein band on membranes was visualized using ECL Western blotting detection reagents (Vazyme) on Bio-Rad ChemiDoc Touch and further quantified by ImageJ software. The primary antibodies were listed as follows: Piezo1 (Alomone Labs, APC-087), APP (Biolegend, SIG-39152), CTFα/β (Biolegend, SIG-39152), GAPDH (Santa Cruz, sc-32233), β-Actin(Santa Cruz, sc-47778).

### Human Aβ_1-42_ ELISA analyses

For Aβ ELISA sample preparation, the cortex and the hippocampus of 5×FAD littermate mice were obtained and sequentially extracted by TBS, TBSX (TBS with 0.5% Triton X-100), and guanidine-HCl (GuHCl) according to Youmans et al. reported, TBS and TBSX fractions represent soluble, and the GuHCl fraction represents insoluble. The Human Aβ1-42 in soluble and insoluble fractions were qualified with Quantikine ™ ELISA kit (R&D Systems) based on the manufacturer’s instructions.

### Aβ Phagocytosis assays

The *in vivo* phagocytosis assay was performed as previously reported with slight modification (Lau et al., 2021). Briefly, *Piezo1*^*fl/fl*^; FAD mice and *Piezo1*^*MGKO*^; FAD at 5 months of age were intraperitoneal with 10 mg/kg methoxy-X04. WT mice were injected with methoxy-X04 to control for methoxy-X04 signal baseline. After anesthesia, mice were perfused with ice-cold HBSS and isolated hippocampi quickly. The isolated hippocampi were minced into pieces, dissociated enzymatically in Papain (5 U/mL, Sigma), DNase I (50 U/mL, Sigma), and Collagenase IV (100 U/mL, Thermo Fisher Scientific) in HBSS (Solarbio) for 30 min at 37°C. Homogenize the dissociated hippocampi by pipetting gently up and down. The homogenates were filtered through a 70-μm cell strainer and centrifuged at 550g for 5 min. Then, hippocampal cells were resuspended in 30% Percoll (Sigma) in ice-cold HBSS and centrifuged at 800 × *g* at 4°C for 15 min without a brake. The cell pellet was resuspended in ice-cold PBS with CD16/CD32 (BD Biosciences) at 4°C for 10 min. After centrifugation and rewash, cells were incubated with CD11b-APC (Invitrogen) and CD45-PE-cy7 (Invitrogen) in fluorescence-activated cell sorting (FACS) buffer (1×PBS containing 1% fetal calf serum, 2 mM EDTA, and 25 mM HEPES) for 45 min at 4°C. Cells were centrifuged and rewashed with FACS buffer for flow cytometry analysis (ACEA Biosciences, Quanteon). For phagocytosis assay *in vitro*, primary microglia cultured on the soft substrate were treated with siRNAs for 36 h, followed by 500 nM FAM-fAβ42 (Anaspec) stimulation. Cells treated with 10 μM cytochalasin D (Calbiochem) for 30 min as negative phagocytosis control. Cells were trypsinized, washed, and incubated with 0.4% trypan blue in FACS buffer for flow cytometry analysis (ACEA Biosciences, Quanteon).

### Microglia isolation from adult mouse brain

Microglia isolation from adult mouse brain was performed as described previously with modifications (Bohlen et al., 2019). Mice were anesthetized and perfused with cold 1×HBSS buffer (Solarbio) quickly. The brain was dissected out and removed olfactory bulb and cerebellum. Obtained brain tissue was placed into a glass homogenizer and homogenized with 1×HBSS (Solarbio) buffer containing DNase I (Sigma, 1:10) and RNasin (Promega, 1:100) on ice gently. Next, tissue homogenates were filtered into a 70-μm cell strainer to a 50-ml conical tube and centrifuged at 550g for 5 min. The supernatants were removed, and pellets were resuspended with 1 ml FACS buffer (1×PBS containing 1% fetal calf serum, 2 mM EDTA, and 25 mM HEPES) with 20 % myelin removal Beads (Miltenyi Biotec) and 1 % RNasin (Promega), and incubated for 15 min at 4 °C. Meanwhile, place a myelin removal column (Miltenyi Biotec) per sample into the magnet separator (Miltenyi Biotec) and rinse with 2 ml FACS buffer. After incubation, dilute suspension to 2 ml FACS buffer and centrifuge at 550 g for 5 min. Then, the suspensions were washed twice and filtered through the myelin removal column (Miltenyi Biotec) in 15-ml conical tubes on ice. The harvested cell suspension was centrifuged at 550g for 5 min. Discard the supernatant and resuspend pellets in the FACS buffer. Cells were incubated with CD16/CD32 (BD Biosciences) blocking antibody to prevent unspecific binding for 10 min at 4 °C. Subsequently, the mixture was centrifuged and washed at 1200 g at 4 °C for 5 min. The cell pellets were resuspended and incubated in FACS buffer with CD11b-APC (Invitrogen) and CD45-PE-cy7(Invitrogen) away from light for 40 min at 4°C. Finally, cells were rewashed and resuspended in FACS buffer containing 5% DNase and 1 % RNasin for flow separation cytometry (MoFlo Astrios EQ 2) to isolate fresh adult microglia.

### RNA isolation, library construction, and bioinformatics analysis

RNA from freshly isolated primary microglia in various groups was extracted with TRIzol reagent (Invitrogen) according to the manufacturer’s instructions. Next, mRNA was purified using poly-T oligo-attached magnetic beads (Vazyme Biotech Co., Ltd). And RNA integrity was assessed using the RNA Nano 6000 Assay Kit of the Bioanalyzer 2100 system (Agilent Technologies, CA, USA). The library fragments (370 ~ 420 bp in length) were processed with the AMPure XP system (Beckman Coulter, Beverly, USA). After the PCR products were assessed using library quality on the Agilent Bioanalyzer 2100 system, the clustering of the index-coded samples was performed on a cBot Cluster Generation System using TruSeq PE Cluster Kit v3-cBot-HS (Illumia). Then, the library preparations were sequenced on an Illumina Novaseq platform, and 150 bp paired-end reads were generated. All sequence reads were trimmed using Cutadapt (version 1.11) and aligned with HISAT2 to Mus musculus GRCm38.After Quality control metrics were finished, expression was performed with RSEM (version 1.2.30). Differentially expressed genes were analyzed by the DESeq2 package in R (Version 3.6.1). The resulting P-values were adjusted using Benjamini and Hochberg’s approach for controlling the false discovery rate. Genes with an adjusted p-value <0.05 found by DESeq2 were assigned as differentially expressed. GO enrichment analyses were performed in the clusterProfiler package (Yu et al., 2012). GSEA analysis was performed using GSEA v2.0.14 software (http://www.broadinstitute.org/gsea/index.jsp). Ingenuity pathway analysis (IPA) was implemented according to Krämer et al. reported (Kramer et al., 2014).

### Behavior tests

For the Morris water maze, Briefly, A circular tank (diameter of 120 cm) filled with opaque water (22°C) was made to perform the Morris water maze. A hidden platform submerged underwater was fixed on the target quadrant. Then, the mice were placed into the maze at one of four random points and performed four training trials per day for five consecutive days. The probe test on the 7^th^ day, the platform was removed, and each mouse placed in the quadrant opposite the platform was allowed 60 s to search the hidden platform. In training and probe trials, mice were tracked by a video camera (Noldus EthoVision XT). Collected data were analyzed by SMART 2.5 VIDEO TRACKING software (Panlab, Harvard Apparatus).

For Y-maze spontaneous alternation analysis, mice were housed and allowed to acclimate in the testing room for 72 h. A Y-shaped maze comprises three identical arms labeled “Unfamiliar arm,” “Initial arm,” and “Familiar arm.” The mice, placed at the end of the “Initial arm,” were allowed to behave in the maze for 5 min freely. On the test period, entries into limbs that pass through all arms were recorded with a video camera (Noldus EthoVision XT) for 10 min. Total arm entries and spontaneous Alternation (%) in 3 different arms were recorded and analyzed.

### Electrophysiological recording

After mice were anesthetized with isoflurane, the brains were dissected out rapidly and placed in ice-cold artificial hypertonic glucose solution (10 mM D-glucose,120 mM sucrose, 10 mM MgSO_4_, 2.5 mM KCl, 1.25 mM NaH_2_PO_4_, 26 mM NaHCO_3_, 0.5 mM CaCl_2_, and 64 mM NaCl). Next, the brains were cut coronally (400 μm thick) by Leica VT1200S vibratome and placed in an artificial hypertonic glucose solution bubbled with carbogen (95% O_2_ + 5% CO_2_). Then, the slices were incubated in ACSF (120 mM NaCl, 1.3 mM MgSO_4_, 3.5 mM KCl, 10 mM D-glucose, 2.5 mM CaCl_2_, 1.25 mM NaH_2_PO_4_, and 26 mM NaHCO_3_), bubbled with carbogen at 32 °C for 1 hour and at RT for at least another 1 hour. The Schaffer collateral inputs to the CA1 region pathway were selected for stimulation with a bipolar stimulating electrode (FHC, Inc.), and the signals were recorded by Multi-Clamp 700B amplifier (Molecular Devices) and digitized using Digidata 1550 A with glass micropipettes (1~3 MΩ) filled with ACSF. Baseline responses were acquired every 20 s with a stimulation intensity that yielded 30% of the maximum response at a frequency of 0.05 Hz. After a 20-min stable baseline recording, LTP was induced by high-frequency stimulation (two trains of 100-Hz stimuli with an interval of 30 s), followed by continued recording for 60 min. The plasticity of synaptic transmission was documented by assessing the initial slope (20~80% rising phase) of field excitatory postsynaptic potentials (fEPSPs).

### Administration of Yoda1

*In vivo* dose of Yoda1 was based on previous reports (Aykut et al., 2020; Romac et al., 2018). Yoda1 (Beidapharma Co., Ltd) was dissolved in 10% DMSO + 90% corn oil. 5×FAD male mice at 1.5 months of age were injected intraperitoneally with Yoda1(3 mg/ kg) every other day until 5.5 months of age, and littermate 5×FAD male mice were treated with DMSO/corn oil (1:9) equivalently.

### Pharmacokinetics of Yoda1

WT mice were intraperitoneal with Yoda1(3 mg/ kg). After that, the whole blood and brain tissues were harvested at 0.5 h, 5 h, and 15 h. Then the whole blood was centrifuged at 4000 rpm for 10 min to collect plasma, and the brain was homogenized. Afterward, 100ul plasma and brain supernatants containing internal standard (IS) [Dooku1(Beidapharma Co., Ltd) (2ng/ml)] were diluted in 400ul methanol (Sigma), followed by ultrasonication thoroughly for 10 min at 4°C. Then, the mixtures were incubated at −20°C for 1 hour and centrifuged at 13000 rpm for 15 min to accelerate protein precipitation. The supernatants were drying at 4 °C using a centrifugal evaporator (Labconco corporation). The samples were detected with an LC-MS system comprised of a Waters Aquity I-class UPLC coupled to a QTRAP 6500+ mass spectrometer (AB SCIEX). Multiple reactions monitoring (MRM) mode was applied to the mass spectrometry. Chromatographic separation was performed using an ACQUITY UPLC BEH C18 column (2.1×50-mm internal diameter; Part No. 186002350, Waters) at a flow rate of 0.40 ml min^−1^ under the condition of 45 degrees column oven. Mobile phase A was 0.1% formic acid(Rhawn); Mobile phase B was methanol. The LC gradient was as follows: 0 to 3 min, 50 to 95% B; 3 to 4min, 95% B; 4 to 4.1 min, 95 to 50% B; 4.1 to 6 min, 50% B.

### Statistical analysis

Statistical analysis was performed with a two-tailed unpaired method (Student’s t-test) for two independent groups and one-way ANOVA for multiple groups with a single variance, all calculated with GraphPad Prism 8. All outcomes presented in this manuscript are shown as mean ± SEM. Significant statistical figure outcomes must obey a p-value upper bound less than 0.05. In addition, the statistical legends marked in the chart are explained as follows: * indicates p < 0.05; ** indicates p < 0.01; *** represents p < 0.005.

## Acknowledgments

We thank Xiang You, Haiping Zheng, Qingfeng Liu, and Xiufeng Sun for their technical support. We thank Dawang Zhou (Xiamen University) for experimental resource. This study was funded by grants from the National Key R&D Program of China (2021YFA1101401 to W.M.); and the Fundamental Research Funds for the Central Universities (20720190086 to W.M.); and the National Natural Science Foundation of China (81602675 to J.H.).

## Author contributions

J.H. and W.M. designed experiments, performed data analyses, and wrote the manuscript. J.H., H.Z., Q.Y., H.S. and G.C. performed experiments. H.Z., B.Z., S. C., Q.C., Z.C. and H.L. performed biochemical analysis of Piezo1-deficient mice. J.H., L.H. and W.M. supervised the study.

---

**Figure S1.**
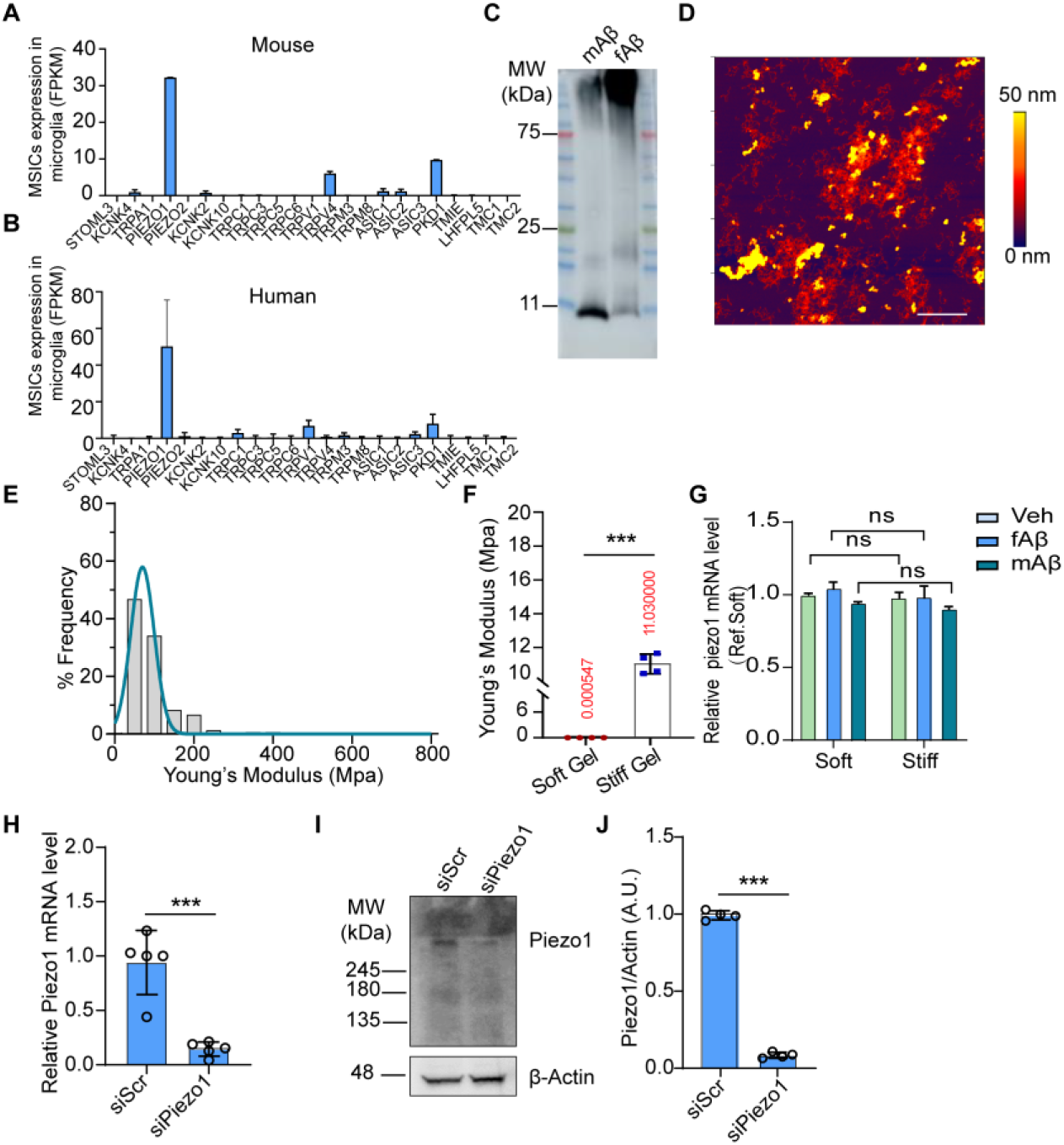
Piezo1 is highly expressed in microglia and fails to be induced by stiffness stimuli at the mRNA level, related to Figure 1. (A and B) Analysis mean FPKM values of various mechanosensory ion channels (MSICs) in mouse microglia (A) from datasets GSE52564 or human microglia (B) from datasets GSE99074. (C) Monomeric Aβ42 peptides (mAβ) and fiberized Aβ42 peptides (fAβ) were immunoblotted with an antibody to detection. (D) AFM image of a fiberized Aβ42 crystal on AFM mica. Scale bar, 1 μm. (E) Frequency distribution of Young’s Modulus (Pa) according to the outcomes of fAβ measured by AFM. (F) Mean shear moduli determined by AFM of our fabricated ‘soft’ and ‘stiff’ hydrogels (n = 4 per group). (G) RT–qPCR analysis of relative Piezo1 mRNA expression level (Ref. Soft hydrogel) on soft and stiff hydrogels for 24 h and cells were simultaneously treated with mAβ (2 μM) or fAβ (2 μM) for 4 h. (H) RT–qPCR analysis of Piezo1 mRNA level in primary microglia transfected with siScr or siPiezo1 for 36 h (n=3 independent experiments). (I and J) Immunoblotting and statistical analysis of Piezo1 levels in primary microglia transfected with siRNAs for 36 h (n=3 independent experiments). Statistics: Mean ± SEM. Unpaired Student’s t-test (F, H, and J); two-way ANOVA with Bonferroni post hoc test (G); ns, no significance; ***p < 0.005.

**Figure S2.**
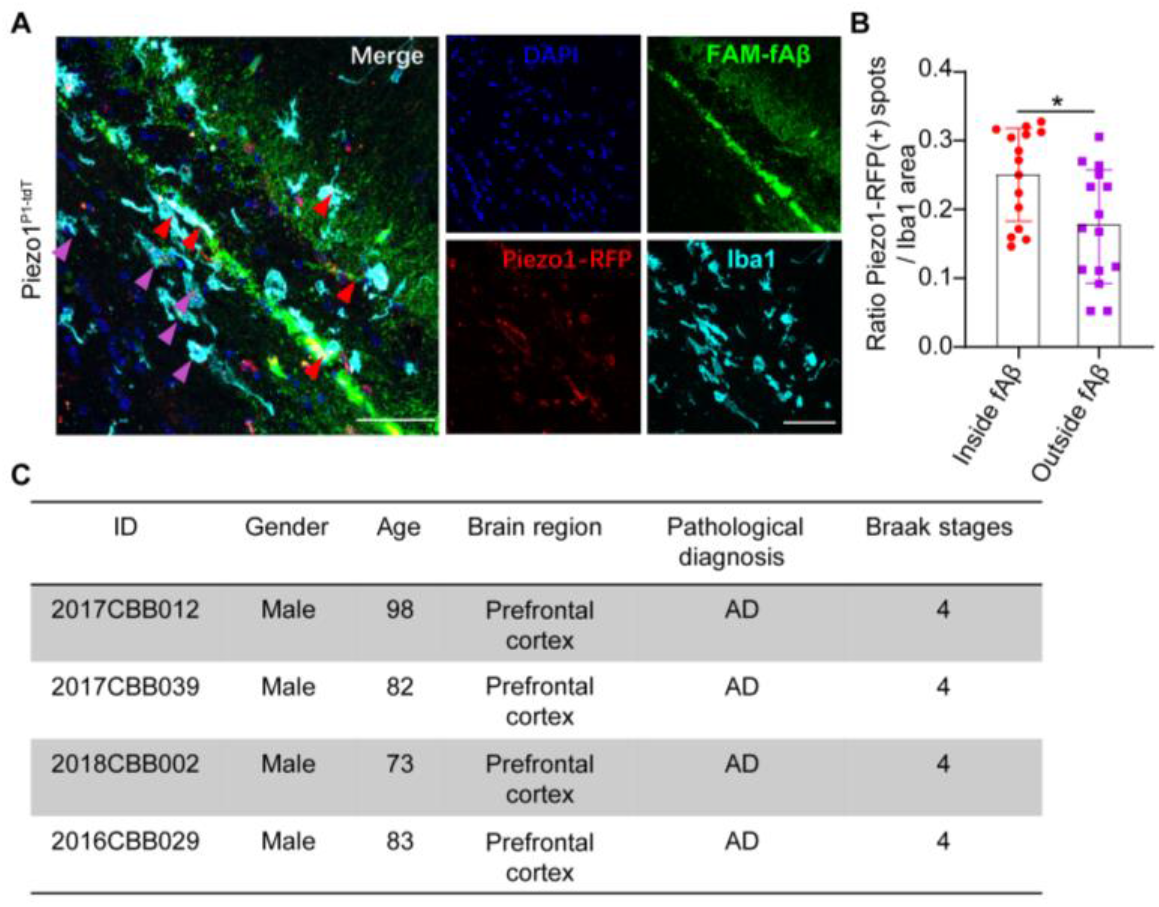
Microglial Piezo1 expression around FAM-fAβ42 deposition in *vivo*, related to Figure 2. (A) Representative image of Piezo1 expression in Piezo1^P1-tdT^ mice at 24 h after FAM-fAβ42 injection. Microglia (Iba1, cerulean blue) outside FAM-fAβ42 (green) were annotated with the purple arrow, and microglia inside FAM-fAβ42 were marked with the red arrow. Scale bar, 50 μm. (B) Statistical analysis of Piezo1 mean fluorescence intensity in Iba1^+^ microglia between the indicated groups (n = 15 independent view fields per group). (C) Detail information of postmortem brain sections from AD patients. Statistics: Mean ± SEM. unpaired Student’s t test (B); *p < 0.05.

**Figure S3.**
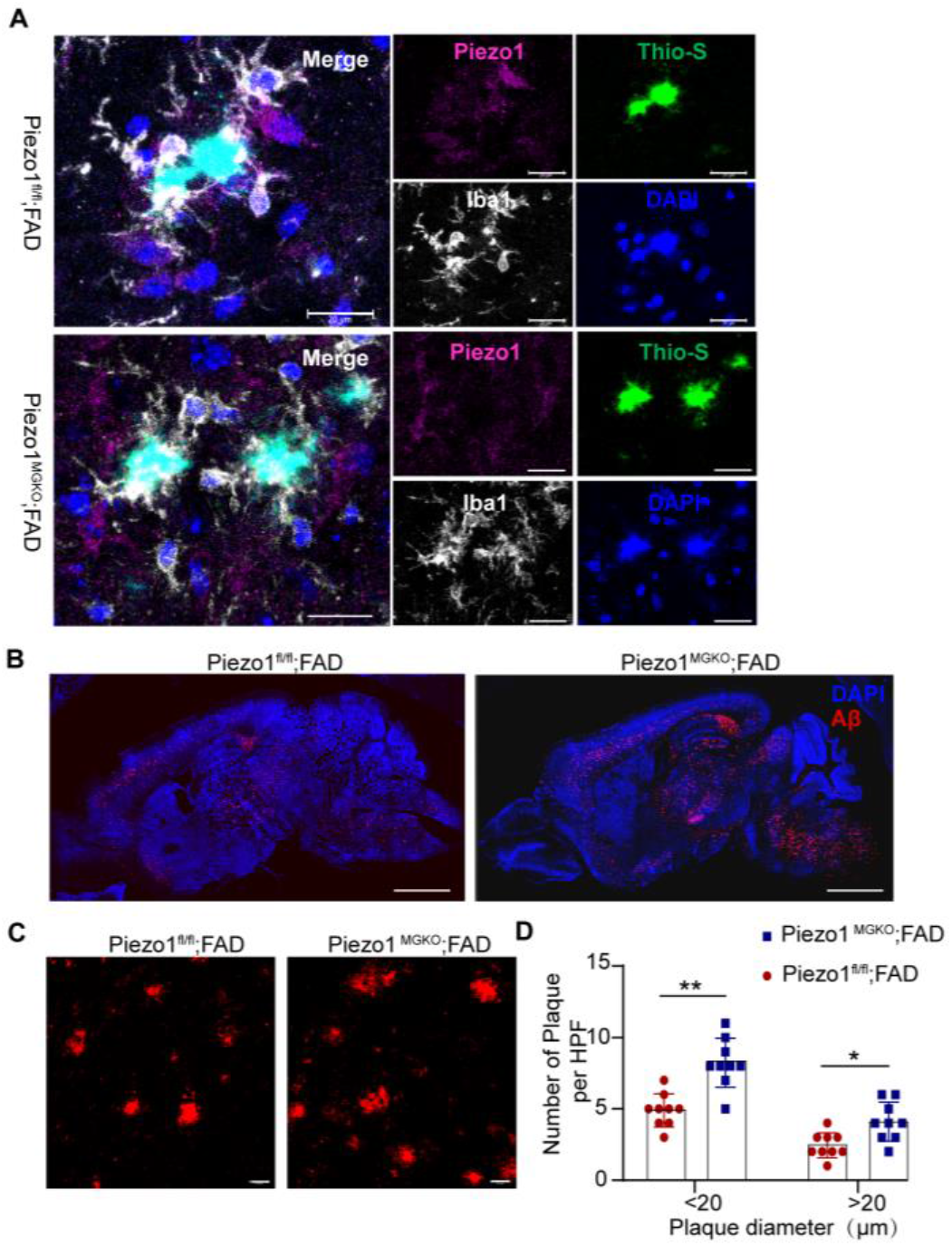
Microglial *Piezo1* deficiency augments Aβ accumulation, related to Figure 3. (A) Representative images of microglia (Iba1, gray), dense-core plaques (Thio-S, green), Piezo1 (purple), and nucleus (DAPI) in indicated groups. Scale bar, 20 μm. (B) Representative sagittal sections from 5-month mouse brains of the indicated groups at least five independent replicates, labeled with Aβ (anti-MOAB2, red). Scale bar, 2 mm. (C) Representative images of Aβ (anti-MOAB2, red) in the hippocampus. Scale bar, 20 μm. (D) Aβ plaques per high power field (HPF) in different genotypes were qualified (n = 9 independent view fields per group). Statistics: Mean ± SEM. unpaired Student’s t-test (D); *p < 0.05; **p < 0.01.

**Figure S4.**
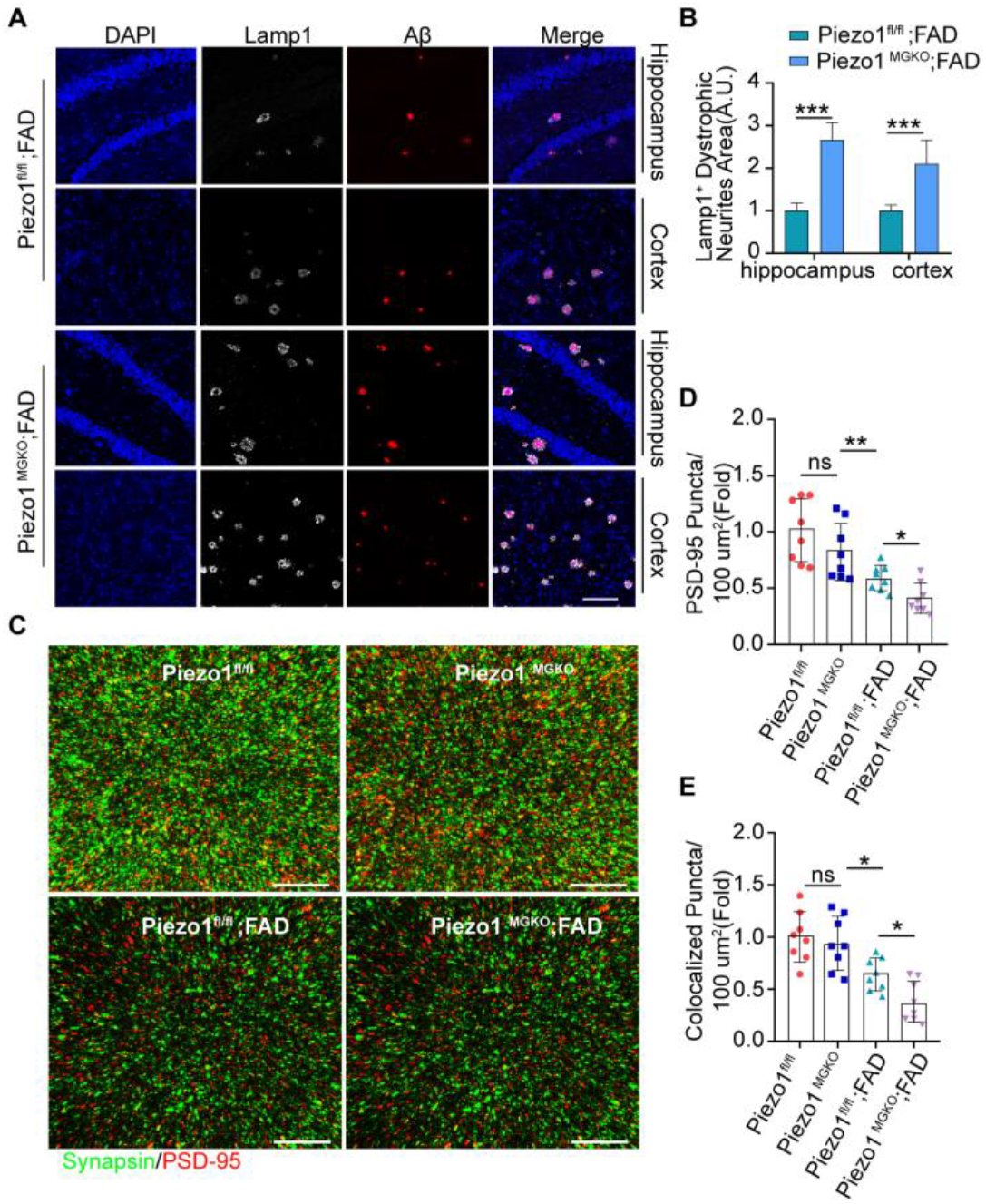
Microglial *Piezo1* deficiency leads to dystrophic neurites and synaptic density reduction, related to Figure 3. (A) Representative images of the holo of Lamp1^+^ dystrophic membranes (white) that surround Aβ plaques (anti-MAOB2, red) in the hippocampus and cortex, respectively., Scale bar, 100 μm. (B) Detail qualification of lamp1^+^ dystrophic neurites area was displayed from both genotypes (n = 3 mice per group). (C) Super-resolution images of synapsin (green) and PSD95 (red) immunoreactive puncta in the hippocampus of the indicated genotypes at 5-month-old. Scale bar, 10 μm. (D and E) Qualification of the relative level of postsynaptic puncta (PSD95 puncta) and colocalized puncta between pre- and postsynaptic puncta in the hippocampus (n = 8 mice per group). Statistics: Mean ± SEM. unpaired Student’s t test (B); one-way ANOVA with Bonferroni post hoc test (D and E); ns, no significance; *p < 0.05; **p < 0.01; ***p < 0.005.

**Figure S5.**
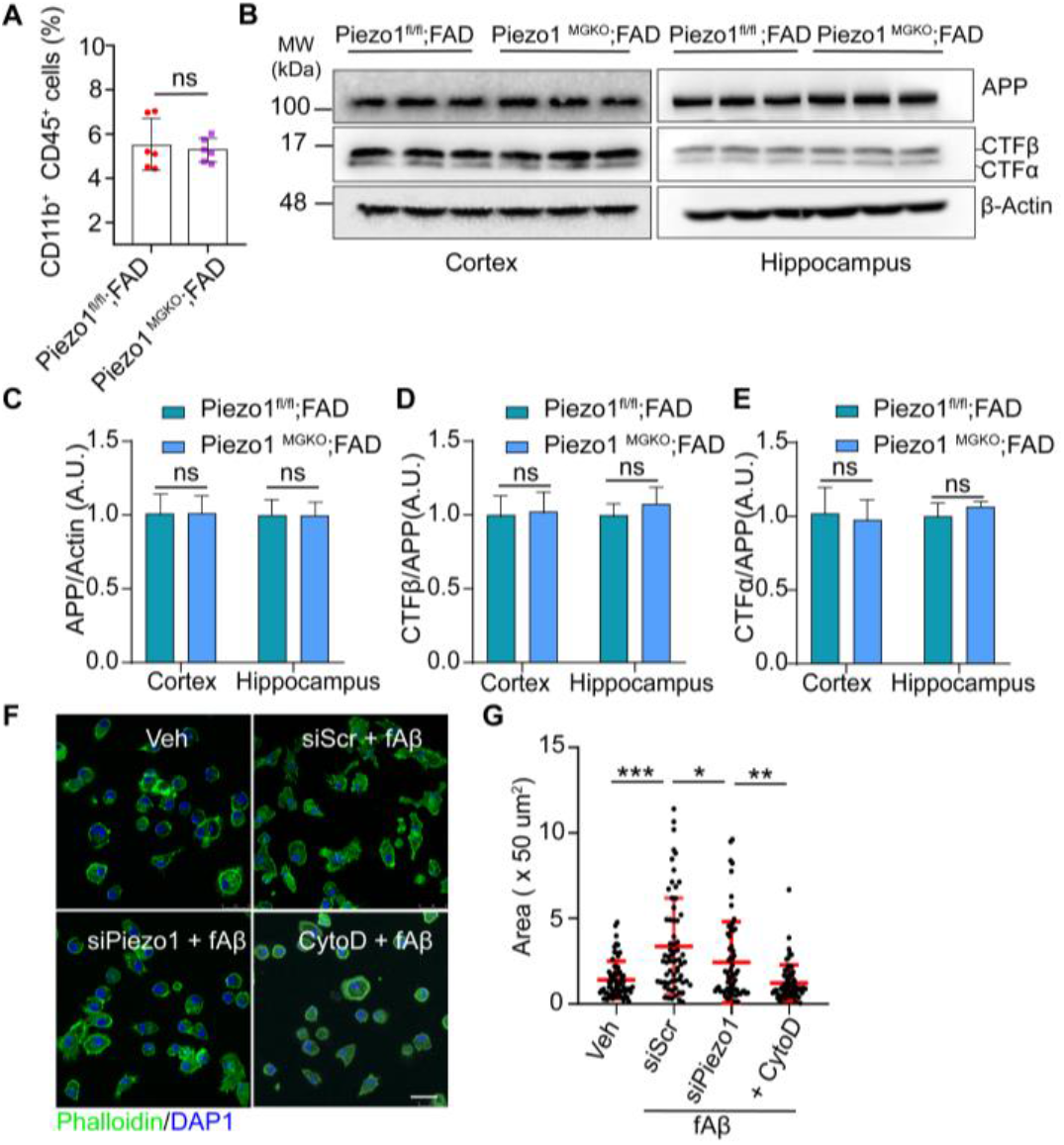
Microglial *Piezo1* deficiency does not influence APP metabolism but relates to phagocytic activity, related to Figure 4. (A) Flow cytometry analysis of the percentage of CD11b^+^ CD45^+^ hippocampal microglial cells between the indicated groups (n = 6 mice per group). (B) Western blotting analysis of APP and its derivates, including APP, β-CTF, and α-CTF in the hippocampus and cortex of the indicated genotypes. (C and E) Quantitation of the protein levels about APP, β-CTF, and α-CTF (n = 3 mice per group). (F) Representative immunofluorescent images of the F-actin (Phalloidin, green). Scale bar, 10 μm. (G) Quantification of the total actin area integrated across primary microglia seeded on the soft gel treated with vehicle, siScr + fAβ, siPiezo1 + fAβ, and fAβ + CytoD, respectively (n= 66 cells in 3 independent experiments per group). Statistics: Mean ± SEM. unpaired Student’s t-test (A, C, D, and E); one-way ANOVA with Bonferroni post hoc test (G); ns, no significance; *p < 0.05; ***p < 0.005.

**Figure S6.**
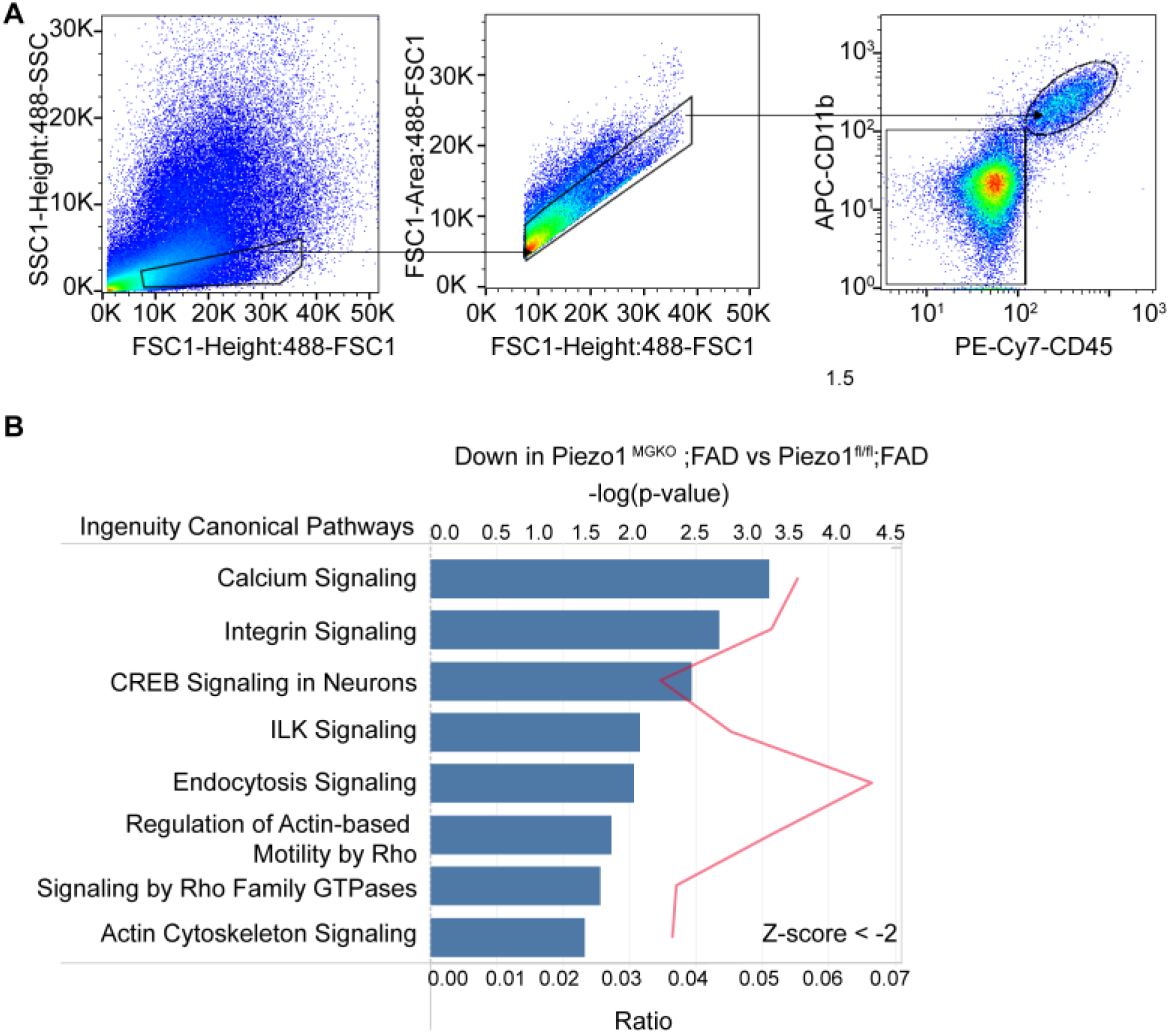
Microglia sorting strategy from adult mice and IPA analysis outcomes between indicated groups, related to Figure 5. (A) The brief FACS sorting strategy for freshly isolated microglia. First, whole cells are gated based on 488-FSC-height/488-SSC-height. In the selected gate, single cells were selected based on 488-FSC-Area/488-FSC-height. Then, fresh microglia were isolated as a PE-Cy7-CD45^+^/APC-CD11b^+^ population. (B) Ingenuity Pathways Analysis (IPA) of Piezo1 ^MGKO^; FAD versus Piezo1^fl/fl^; FAD revealed that microglial Piezo1 deficiency was associated with decreased expression of genes relating to integrin, endocytosis, and actin cytoskeleton-based signaling pathways, etc.

**Figure S7.**
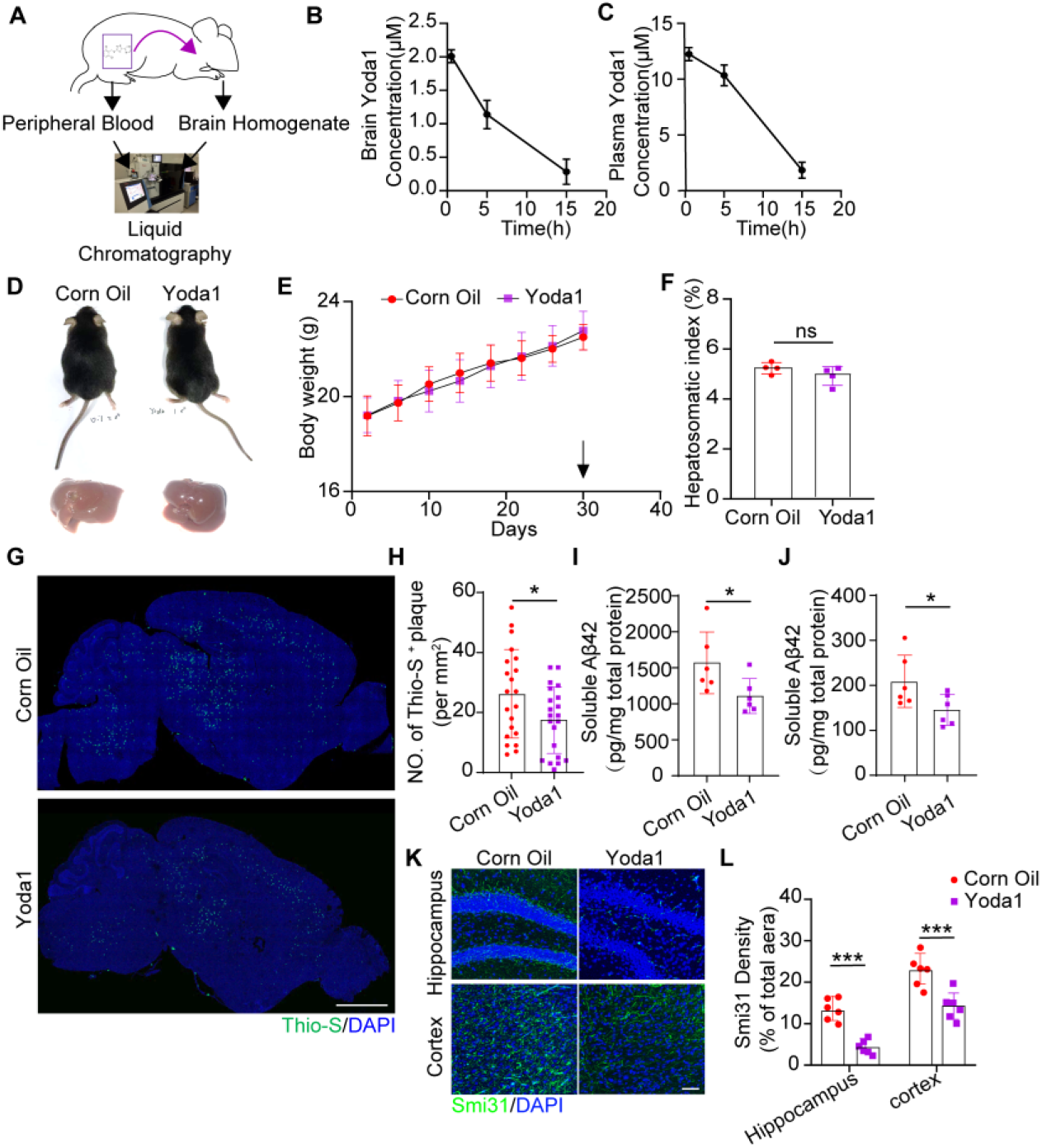
Pharmacokinetic properties of Yoda1 and its protective effects on 5×FAD mice, related to Figure 6. (A) Brief schematic of the methodology for pharmacokinetics of Yoda1. (B and C) Brain and plasma concentrations of Yoda1 (μM) were detected with LC-MS/MS after intraperitoneal administration at body weight (3 mg kg^−1^) (n = 3 mice per time point). (D) Body phenotype and liver size of 5-month-old 5×FAD mice treated with Corn Oil or Yoda1. (E) Dynamic monitor of body weight (g) for 30 days before the end of the last day of medication. (F) The liver weight (g) / body weight was calculated for the hepatosomatic index after drug administration. (G) Representative images of sagittal brain sections from 5-month-old indicated 5×FAD mice treated for staining dense-core plaques (Thio-S, green). Scale bar,1 mm. (H) No. of Thio-S^+^ plaque (per mm^2^) was qualified (n = 21 independent view fields per group). (I and J) Quantitative analysis of the relative content level of soluble (I) and insoluble (J) Aβ42 in hippocampus measured by ELISA (n = 6 mice per group). (K) Immunofluorescent staining of Smi31 in cortex and hippocampus. Scale bar, 50 μm. (L) Statistical analysis of Smi31 density of 5×FAD mice treated with Corn Oil or Yoda1 (n = 6 independent view fields per group). Statistics: Mean ± SEM. unpaired Student’s t test (F, H, I, J, and L); ns, no significance; *p < 0.05; ***p < 0.005.

